# An N-Terminally Acetylated Degron Specifically Recognized by the CTLH^MKLN1-FAM72A^ E3 Ubiquitin Ligase for Regulatory Protein Degradation

**DOI:** 10.64898/2026.09.19.752822

**Authors:** Maosheng Yang, Shuqi Yao, Zheng Fang, Wenjie Ren, Yanyang Yang, Qizhou Song, Wen Yang, Hui Wang

## Abstract

N-terminal acetylation of cellular proteins has been implicated in their proteasomal degradation through the Ac/N-degron pathway. Despite representing a major branch of the N-degron system, only a limited number of degrons and their motifs have been identified and characterized in this pathway, and the mechanism underpinning the recognition of N-terminal acetylation has remained largely elusive. Here, by interrogating FAM72A-mediated degradation of UNG2, we pinpointed the N-terminally acetylated sequence of UNG2 as a bona fide Ac/N-degron specifically recognized by the CTLH^MKLN1-FAM72A^ E3 ubiquitin ligase. Rather than relying solely on a single acetylated N-terminal residue, this degron features a consensus Nt-AcM_1_-[ILV]_2_-G_3_ motif that governs both E3 targeting and the intracellular stability of the neo-substrates uncovered through proteomic screenings. Cryo-EM structure of the UNG2-FAM72A-MKLN1 complex further elucidated the molecular details supporting the high specificity of degron recognition. Notably, this degron pathway is subject to multilayered regulation, highlighting the characteristic conditionality of degron system. Our findings thus define a distinct AcM-[ILV]-G/N-degron pathway and provide mechanistic insights into degron recognition dictated by both N-terminal acetylation and sequence motif context.

## Introduction

As a critical arm of proteome maintenance in eukaryotic cells, protein degradation primarily relies on the Ubiquitin-Proteasome System (UPS) for selective, rapid protein turnovers, and the Autophagy-Lysosome Pathway (ALP) to handle bulk or organelle-level clearance (Pohl & Dikic, 2019). To ensure precise timing and specificity in protein breakdowns, both pathways, particularly the UPS, are driven by the recognition between certain cellular machinery such as E3 ubiquitin ligases, and degrons, minimal intrinsic element within proteins to signal their degradation (Ravid & Hochstrasser, 2008; Sherpa *et al*, 2022; Zhang *et al*, 2025). Initially identified at the N-termini of proteins as “destabilizing” residues in yeast (Bachmair *et al*, 1986), degrons were first characterized to be short linear sequence motifs that mediate interactions with E3 ligases or other degron-recognizing factors (Varshavsky, 1991). Over subsequent decades, as additional degradation signals and their recognition mechanisms were uncovered, degrons have come to be understood as conditional and context-dependent rather than static. They are often regulated by protein folding state, complex assembly, post-translational modifications (PTMs), proteolytic processing, or ligand binding (Carrillo Roas *et al*, 2025; Gray *et al*, 2001; Guharoy *et al*, 2016; Ito *et al*, 2010; Ivan *et al*, 2001; Jaakkola *et al*, 2001; Mark *et al*, 2023; Piatkov *et al*, 2012; Ravid & Hochstrasser, 2008; Shemorry *et al*, 2013; Tan *et al*, 2007; Winston *et al*, 1999; Yagita *et al*, 2023), becoming active under specific physiological or biochemical conditions.

Degrons can be classified as internal or terminal based on their position within the protein sequence. Internal degrons typically comprise longer and more complex motifs and are subject to tight control in key signaling pathways (Zhang *et al*., 2025; Zhang *et al*, 2023). By contrast, terminal degrons lean on the identity and modification of residues at the extreme N- or C-ends of proteins to determine their half-life, influencing processes such as protein quality control, metabolism, stress responses, and development (Sherpa *et al*., 2022; Varshavsky, 2019). Established as a prototype of degron-dependent degradation, the N-degron system encompasses several canonical branches, including the Arg/N-degron, Ac/N-degron, Pro/N-degron, and Gly/N-degron pathways (Varshavsky, 2024). These pathways operate through dedicated E3 ubiquitin ligases, such as the UBR family, MARCH6 and Not4, Gid4 in the yeast GID (glucose-induced degradation) complex, and the CRL2 (Cul2-RING ubiquitin ligase) complex substrate receptors ZYG11B and ZER1 (Chen *et al*, 2017; Hwang *et al*, 2010; Shemorry *et al*., 2013; Tasaki *et al*, 2005; Timms *et al*, 2019). Concurrently, alongside the integration of systems biology and proteomic studies, the degron landscape has been expanded to include the more recently defined C-degron pathways, which are primarily mediated by Kelch-repeat and FEM1 family substrate receptors of the CRL2 complex and certain substrate receptors of the CRL4 complex (Sherpa *et al*., 2022; Zhang *et al*., 2025). It is of note that, beyond being intrinsically encoded, terminal degrons can be created or activated by PTMs such as arginylation, acetylation, deamination, and oxidation happened at the N-termini of proteins, as well as amidation and cyclization of glutamine or asparagine at the C-termini (Ichikawa *et al*, 2022; Muhar *et al*, 2025; Varshavsky, 2024). Meanwhile, proteolytic cleavage can give rise to new terminal degrons by exposing neo-protein ends at the cleavage site (Lin *et al*, 2018; Ravalin *et al*, 2019).

Recognition of terminal degrons is executed by the substrate receptor domains within E3 ubiquitin ligases or their associated complexes. A handful of structural studies have demonstrated the diversity of this recognition, showing that these domains adopt distinct folds capable of engaging one or multiple cognate degrons (Sherpa *et al*., 2022). For example, Arg/N-degrons are recognized by UBR-box domains in UBR family E3 ligases (Choi *et al*, 2010; Matta-Camacho *et al*, 2010); C- and Pro/N-degrons by β-barrel folds in FBXO31 and GID substrate receptors (Dong *et al*, 2018; Li *et al*, 2018; Shin *et al*, 2021); diGly/C-degrons by the Kelch-type β-propeller domains in CRL2 substrate receptors (Rusnac *et al*, 2018); and Gly/N- and Arg/C-degrons by helical repeats in CRL2 substrate receptors (Chen *et al*, 2021; Yan *et al*, 2021a; Yan *et al*, 2021b). However, despite decades of extensive research, the number of identified degrons and degron motifs, particularly those contextually created or activated, and their corresponding recognins remains relatively limited, and the molecular basis of their recognition are even less well-understood.

Among the known E3 ubiquitin ligases committed to degron-dependent degradation, the human CTLH complex, named for the shared ‘C-terminal to LisH (CTLH)’ domain present in most of its subunits, is an evolutionary conserved counterpart of the yeast GID complex (Francis *et al*, 2013; Kobayashi *et al*, 2007; Santt *et al*, 2008). It functions as a multiprotein RING-type E3 ligase and operates across a broad range of biological processes encompassing metabolic regulation, cell signaling, and development, to remodel the proteome in response to cellular cues (Maitland *et al*, 2022). As a modular architecture, the CTLH E3 complex is organized around a central scaffold composed of RANBP9 (or RANBP10) and TWA1, and a catalytic module formed by a heterodimer of MAEA and RMND5A (or its paralog RMND5B). This core is complemented by interchangeable and combinatorial substrate receptors and their associated adaptors including WDR26, muskelin (MKLN1), GID4, ARMC8, and FAM72A (Alpi *et al*, 2025). The incorporation of these variable subunits enables the CTLH E3 complex to form distinct higher-order assemblies, facilitating the efficient recognition and capture of oligomeric substrates (Mohamed *et al*, 2021; Sherpa *et al*, 2021; van Gen Hassend *et al*, 2023). Thus far, most of the substrate receptors described above have been functionally and/or mechanistically linked to specific degron pathways. First, similar to its yeast ortholog, human GID4, the canonical substrate receptor teaming up with its adaptor ARMC8, is able to recognize Pro/N-degrons that are often conditionally generated or activated (Owens *et al*, 2024; Yi *et al*, 2024). In addition, WDR26 is primarily involved in an internal basic degron pathway by recognizing a consensus sequence of K-R-x-Φ (where Φ denotes a bulky hydrophobic residue) (Gottemukkala *et al*, 2024). Lastly, MKLN1 can function as a substrate receptor to promote DR-like C-degron dependent degradation (Chen *et al*, 2026; Grant *et al*, 2026).

In contrast, FAM72A, despite its recent identified partnership with MKLN1 as the substrate receptor to degrade Uracil-DNA Glycosylase2 (UNG2) (Barbulescu *et al*, 2024), an enzyme that initiates the base excision DNA repair pathway, and thereby contributions in antibody diversification and cancer progression (Feng *et al*, 2025; Feng *et al*, 2021; Rogier *et al*, 2021), has not yet been connected to any known degrons or related pathways. We herein investigated the recognition of UNG2 by FAM72A and found that both this interaction and the subsequent FAM72A-regulated polyubiquitination of UNG2 strictly depend on N-terminal acetylation of UNG2. By means of integrated biochemical, structural, quantitative proteomic, protein interactomic, and cellular analyses, we identified the Nt-acetylated sequence of UNG2 as a bona fide Ac/N-degron defined by a consensus motif of Nt-AcM_1_-[ILV]_2_-G_3_. This degron mediates the stability of a group of biologically important neo-substrates through specific recognition by the substrate receptor FAM72A and its associated CTLH E3 ligase. In accordance with the conditional nature of degron-dependent degradation, we further demonstrated that the AcM-[ILV]-G/N-degron pathway is regulated at multiple levels, including N-terminal acetylation status, substrate receptor abundance, and the functional context of its substrates.

## Results

### N-terminal acetylation of UNG2 dictates its targeting by the CTLH^MKLN1-FAM72A^ E3 ligase

To probe the recognition of UNG2 by FAM72A, we reconstituted their interaction in a bacterial recombinant purified system using pull-down assays. Compared with conventionally N-tagged construct, only C-tagged UNG2 was recognized by FAM72A (Fig. 1A). Further mapping with a series of C-tagged UNG2 truncation constructs designed according to domain organization and sequence conservation (Fig. EV1A,B) revealed that the N-terminal region of UNG2 containing PCNA binding site is required for the FAM72A engagement (Fig. 1A), consistent with the FAM72A-binding site (first 25 amino acids of UNG2) identified in previous cell-based studies (Guo *et al*, 2008). To isolate the UNG2-FAM72A complex, purified C-tagged UNG2 and FAM72A were mixed at a 1:1 molar ratio and applied to size exclusion chromatography. To our surprise, only a small fraction of the proteins formed a stable complex, whereas the majority were eluted separately (Fig. 1B). To test whether partial N-terminal degradation of UNG2 compromised its recognition by FAM72A, we generated an N-terminal SUMO-tagged UNG2 construct that could be cleaved by ULP1 to release a fresh, intact N-terminus for recognition prior to the pull-down assay. Unexpectedly, this construct completely lost the ability to interact with FAM72A (Fig. 1A). These findings therefore suggest that the N-terminus of UNG2 undergoes a low-frequency processing event that is nevertheless critical for its recognition by FAM72A.

**Figure 1.**
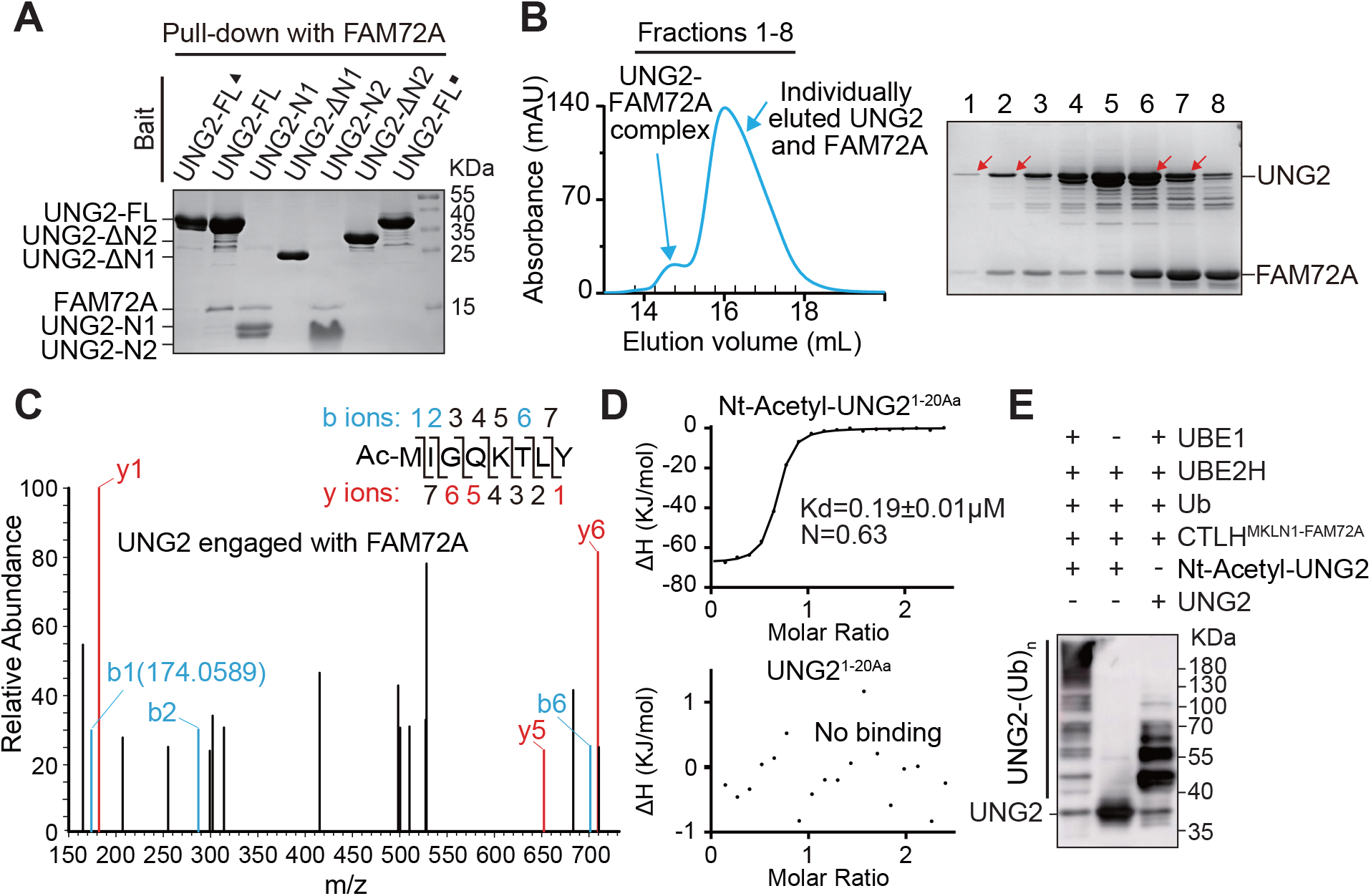
N-terminal acetylation of UNG2 is essential for recognition and ubiquitination by the CTLH^MKLN1-FAM72A^ E3 ligase. (A) Nickel pull-down assay using purified FAM72A and purified C-terminal His-tagged UNG2 constructs spanning different regions. The black square indicates the UNG2 construct with SUMO protease cleavage. As a control, an N-terminal His-tagged UNG2 construct was included and is indicated by a black triangle. (B) Size-exclusion chromatography analysis of the UNG2-FAM72A interaction. Eluted fractions were resolved by SDS-PAGE and analyzed by Coomassie staining. Protein bands subjected to LC-MS/MS analysis are indicated by red arrows. (C) Representative MS/MS fragmentation spectrum of an N-terminally acetylated peptide from UNG2 bound to FAM72A. The sample was obtained from the protein bands indicated by red arrows in (B) (lanes 1 and 2). The b and y ions represent the two complementary peptide fragments generated by higher-energy collision dissociation, and the corresponding product ion peaks are labeled. The observed *m/z* value of the b1 ion is consistent with N-terminal acetylation of the methionine residue. (D) ITC binding curves for FAM72A titrated with UNG2 peptide (1-20Aa) with or without N-terminal acetylation. The corresponding peptides and their binding affinities are indicated. (E) *In vitro* ubiquitination of full-length UNG2 with or without N-terminal acetylation by the CTLH^MKLN1-FAM72A^ E3 complex. Ubiquitination was detected by immunoblotting with a UNG2-specific antibody.

To unveil the nature of this processing event, protein samples from the stable complex (Fig. 1B, lanes 1 and 2) and individually eluted UNG2 in the subsequent peak (Fig. 1B, lanes 6 and 7) were respectively subjected to mass spectrometry (MS) analysis. Tandem mass spectrometry (MS/MS) revealed that UNG2 in complex with FAM72A was N-terminally acetylated at Methionine 1 (M1), whereas UNG2 incapable of interacting with FAM72A consisted of a heterogeneous population, including non-modified, M1-cleaved, M1-methylated, and M1-acetylated forms (Figs. 1C and EV1C; Dataset EV1). To further validate the role of N-terminal acetylation in UNG2 recognition by FAM72A, we synthesized peptides containing residues 1-20 of UNG2, with or without M1 acetylation, based on our biochemical mapping and sequence conservation analyses (Fig. EV1B). Their binding affinities to FAM72A were quantitatively measured by isothermal titration calorimetry (ITC). In line with our previous observations, the N-terminally acetylated peptide bound FAM72A with high affinity (∼190nM), whereas the unmodified WT peptide showed no detectable binding (Fig. 1D).

Considering that N-terminal acetylation is far less prevalent in bacteria than in eukaryotes (Schmidt *et al*, 2016), we sought to recapitulate the N-terminal acetylation-mediated UNG2-FAM72A interaction in mammalian cells. Purified FAM72A was immobilized as bait for affinity pull-down assays against HEK293T cell lysate (Fig. EV1D), and the distinctly captured UNG2 band was subjected to MS analysis. Consistent with our findings in bacterial system, FAM72A-complexed UNG2 was unequivocally N-terminally acetylated (Fig. EV1E; Dataset EV1). In addition to UNG2, several distinct bands corresponding to the UNG2-associated RPA complex and the CTLH E3 ligase component MKLN1 were also detected on the gel (Fig. EV1D).

Subsequent MS analyses identified other components of the CTLH E3 complex within the FAM72A-interacting proteome (Fig. EV1F; Dataset EV2), in agreement with previous report that FAM72A recruits the CTLH E3 complex for UNG2 degradation (Barbulescu *et al*., 2024). We next examined the effect of UNG2 N-terminal acetylation on its ubiquitination by the CTLH^MKLN1-FAM72A^ E3 ligase. *In vitro* ubiquitination assays showed that N-terminally acetylated UNG2 was efficiently polyubiquitinated by the assembled E3 complex while unmodified UNG2 displayed substantially reduced polyubiquitination (Fig. 1E). Taken together, we biochemically demonstrate that UNG2 recognition and ubiquitination by the CTLH^MKLN1-FAM72A^ E3 ligase are determined by its N-terminal acetylation.

### Molecular basis of UNG2-FAM72A recognition mediated by N-terminal acetylation

To understand how N-terminal acetylation of UNG2 mechanistically regulates its interaction with FAM72A in the context of the CTLH E3 ligase, we reconstituted the UNG2-FAM72A-MKLN1-RANBP9-TWA1 complex and analyzed its structure using single-particle cryo-electron microscopy (cryo-EM) (Fig. EV2). A final structural model containing UNG2, FAM72A, and MKLN1 was obtained at a resolution of 2.81Å through a combination of AlphaFold-assisted model fitting, manual rebuilding, and refinement (Figs. 2A,B and EV2; Table EV1). Overall, the UNG2-FAM72A-MKLN1 complex adopts a symmetric dimeric architecture with FAM72A bridging UNG2 and MKLN1 in each monomer, which is in line with the previously proposed organization of this trimeric module within the ring-shaped CTLH E3 ligase (Barbulescu *et al*., 2024). The discoidin, LisH, CTLH, and CRA^C^ domains of MKLN1 are fully resolved in our model, but the kelch and part of CRA^N^ domains are absent due to their flexibility or lack of binding partner. The LisH, CTLH, and looped-back CRA^C^ domains form an intertwined α-helical structural module that facilitates MKLN1 homodimerization. The barrel-shaped discoidin domain of MKLN1 utilizes the top with protruding loops to engage FAM72A, linking it to the CTLH E3 assembly (Fig. 2A,B).

**Figure 2.**
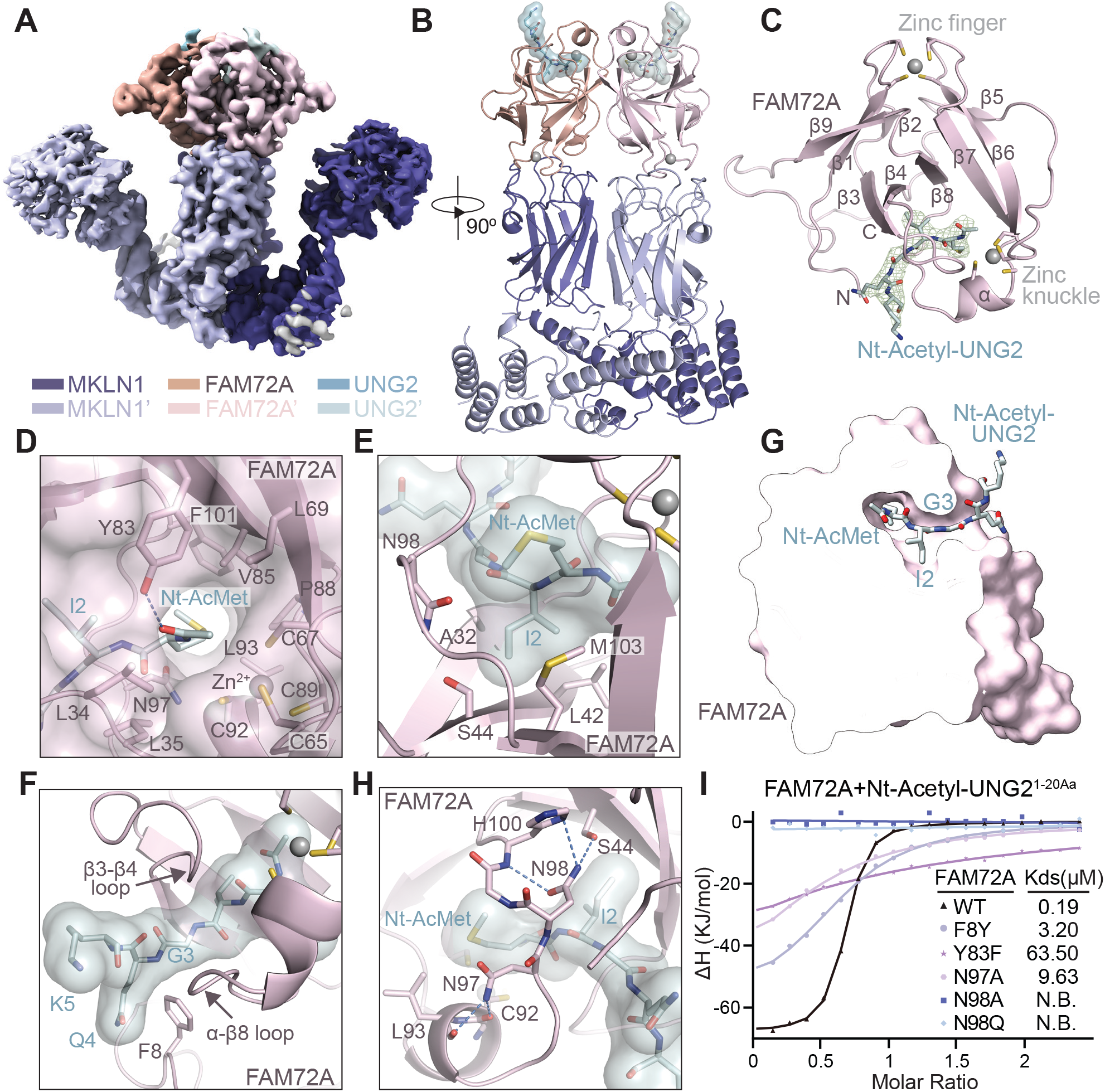
Cryo-EM structure revealing recognition of N-terminally acetylated UNG2 by FAM72A. (A) C2-symmetrized cryo-EM map of the UNG2-FAM72A-MKLN1 complex revealing its dimeric architecture. MKLN1 is shown in slate blue, FAM72A in pink, and UNG2 in teal. Different shades of each color distinguish the two monomers of each component. (B) Overall structure of the UNG2-FAM72A-MKLN1 complex shown in an orthogonal view to (A). The FAM72A-MKLN1 subcomplex is displayed as a ribbon representation using the same color scheme as in (A), while the N-terminally acetylated peptide of UNG2 is shown as sticks with a transparent surface. Zinc ions are shown as gray spheres. (C) Structure of FAM72A in complex with the N-terminally acetylated peptide of UNG2. FAM72A is shown in pink ribbon diagram, and Nt-Acetyl-UNG2 is shown as teal sticks overlaid with a mesh electron density map contoured at 16 σ. The nine β-strands are labeled sequentially, with the N and C termini of the protein indicated. Zinc ions are shown as gray spheres and their cysteine ligands as sticks. (D-F) Close-up views of the selective recognition of Nt-Acetyl-UNG2 by FAM72A, featuring the Nt-AcM1 residue (D), I2 residue (E), and G3 residue (F). Nt-Acetyl-UNG2 is shown as teal sticks with or without a surface, and FAM72A is shown as pink ribbon with or without a surface. Selected interface residues are displayed as sticks. Blue dashed lines indicate hydrogen bonds, and zinc ions are shown as gray spheres. In (F), the β3-β4 and α-β8 loops of FAM72A are indicated. (G) Clipped surface view of FAM72A highlighting the tunnel, cavity, and bottleneck that accommodate the Nt-AcM1, I2, and G3 residues of Nt-Acetyl-UNG2, respectively. The Nt-Acetyl-UNG2 peptide is shown as teal sticks. (H) Roles of FAM72A residues N97 and N98 in stabilizing both Nt-Acetyl-UNG2 and the pocket-shaping loops of FAM72A through an intramolecular interaction network. FAM72A is shown as pink ribbon, and Nt-Acetyl-UNG2 is shown as teal sticks with a surface. Selected residues are displayed as sticks, and dashed lines indicate hydrogen bonds and polar interactions within the network. (I) ITC binding curves for WT FAM72A and its variants titrated with the Nt-Acetyl-UNG2 peptide (1-20Aa). The corresponding FAM72A variants and their binding affinities are indicated. N.B., no detectable binding.

Similar to canonical CULT (Cereblon domain of Unknown activity, binding cellular Ligands and Thalidomide) domains, monomeric FAM72A adopts a β-tent fold, consisting of two sets of β-meanders arranged at roughly right angles and pinned together at their tip by a zinc ion coordinated by four cysteines (residues C18, C21, C74, and C77), featuring C-X_2_-C spacing motif (Figs. 2C and EV3A-D). This zinc ion and its coordinating loops directly interact with the discoidin domain of MKLN1, forming an interface between FAM72A and MKLN1. A β-hairpin insertion between strands β2 and β5 penetrates through the molecule toward the base of the tent. This hairpin, along with its hairpin loop and the loops connecting β5 to β6 and β7 to β8 (a short α-helix included), forges a cradle-shaped pocket that accommodates the binding ligand (Fig. 2C). Unlike canonical CULT domains, FAM72A lacks the typical aromatic-cage that facilitates ligand binding (Lupas *et al*, 2015). Instead, its binding pocket harbors a second zinc ion coordinated in distinct C-X_1_-C configuration by residues C65 and C67 within the β5-β6 loop (Figs. 2C and EV2H, 3D). The constrained coordination geometry forces a pinched conformation of this zinc knuckle, in combination with the adjacent ligated short α-helix, jointly creating a hydrophobic surface within the pocket, where sits the N-terminally acetylated pentapeptide of UNG2 (Fig. 2C).

According to the density captured in the ligand-binding pocket, five amino acids at the N-terminus of UNG2, starting from the acetylated M1 residue, were clearly modeled, leaving the majority of UNG2 remained unresolved due to the lack of density (Fig. 2C). A close examination of the pentapeptide binding area revealed that the first three N-terminal residues of UNG2 are specifically recognized by FAM72A primarily through hydrophobic interactions (Fig. EV3D).

For the N-acetyl-methionine at position 1 (Nt-AcM1), the methyl group of the acetyl moiety leans against the hydrophobic surface generated by the pinched zinc knuckle, while the carbonyl oxygen is anchored by a hydrogen bond with the hydroxyl group of Y83 (Fig. 2D). The side chain of M1 also packs against the same hydrophobic surface. Together with the acetyl group, it lies snugly in a hydrophobic tunnel traversing the binding pocket, which is maintained by residues L34, L35, L69, V85, P88, L93, and F101, along with the side-chain carbon backbone of N97 (Fig. 2D). Isoleucine at position 2 (I2) is secured within a fitted hydrophobic cavity composed of residues A32, L42, and M103, as well as the side-chain carbon backbones of S44 and N98 (Fig. 2E). The third residue glycine (G3) is hydrophobically stabilized by F8, whose aromatic ring is oriented toward the backbone of G3. In particular, G3 is firmly clamped by both the β-hairpin insertion and the α-β8 loop (Fig. 2F), leaving limited space around the residue backbone that would accommodate either a very small side chain or none at all. This spatial constraint, evident from the clipped surface representation, forms a bottleneck within the peptide-binding pocket and highlights the stringency of UNG2-FAM72A recognition (Fig. 2G). Beyond the first three residues, glutamine at position 4 (Q4), located at the entrance of the binding pocket, makes a loose hydrophobic contact with the side chain of F8, while lysine at position 5 (K5) stays outside the binding pocket and shows no detectable interaction with FAM72A (Fig. 2F).

Besides the residue-specific interactions, FAM72A residues V33 and L35 in the β-hairpin loop, along with residues C96 and N98 in the α-β8 loop, engage the peptide backbone through main chain-mediated contacts, reinforcing peptide recognition via an extensive hydrogen bond network (Fig. EV3E). Notably, within this network, both the carbonyl and amide groups of peptide residue G3 are gripped by FAM72A residue V33, effectively locking the position of G3 in place. Among all FAM72A residues participating in peptide engagement, N97 and N98 emerge as essential organizational elements. These residues not only contribute to the recognition of Nt-AcM1 and I2, as well as to the supportive hydrogen bond network, but also dictate the local conformation of the pocket-shaping loops. Specifically, their side chains establish multiple polar interactions with residues in both the β-hairpin and the α-β8 loop, stabilizing an intramolecular configuration that is favorable for peptide binding (Fig. 2H).

We next tested the functional importance of the key FAM72A residues identified by structural analyses in mediating UNG2 recognition. A series of FAM72A mutants were expressed and purified, and their binding affinities toward a synthesized Nt-acetylated UNG2 peptide were quantified by ITC experiments. As expected, substitution of Y83, which recognizes the acetyl group, dramatically weakened the binding affinity to approximately 60µM, while disruption of the hydrophobic stabilization surrounding G3 (F8Y) reduced the binding by more than 15-fold (Fig. 2I). Remarkably, mutations affecting the critical organizational residues N97 and N98 (N97A, N98A, and N98Q) substantially impaired the interaction if not completely abolished the binding, underscoring their indispensable roles in UNG2 recognition (Fig. 2I).

Collectively, these structural and biochemical analyses unfold the molecular basis of UNG2-FAM72A recognition, which relies on both the Nt-acetylated methionine and the structural stringency imposed on the subsequent peptide residues within the binding pocket.

### FAM72A undergoes dimerization and stability regulation mediated by the CTLH E3 ligase

Expanding our analysis to the entire assembly, we examined the symmetric dimeric structure of the UNG2-FAM72A-MKLN1 complex, which is primarily driven by MKLN1 homodimerization via its LisH and CRA^C^ domains at the base of the architecture (Fig. 2A,B). The LisH domain contains a long α-helix (αL) that steadily holds up the discoidin domain of MKLN1, followed by a short α-helix (αS). Both helices participate in dimer formation as previously reported (Delto *et al*, 2015). On top of that, residues L729 and L732 in the looped-back CRA^C^ domain of each monomer slide into the hydrophobic core like two bolts, further securing the homodimerization of MKLN1 (Fig. EV4A).

Interestingly, FAM72A also forms a dimer within the assembly, mainly through a surface located on the backside of the ligand-binding pocket. This surface comprises the β5-β8 β-meander and connecting loops, as well as the C-terminal region captured in our model (Fig. 3A). The dimer interface of FAM72A is considerably extensive, encompassing nearly half of the molecule surface, and is predominantly maintained by a hydrophobic core formed by residues such as I47, P48, T50, F55, C59, F61, H84, I86, V87, P88, L93, F101, W102, and I131 (Figs. 3A and EV3D). Surrounding the core, specific polar interactions involving residues K68 and D46, along with E130 from the C-terminus, bridge the monomers to anchor the orientation in place. To determine whether the observed dimerization occurs under native conditions, purified FAM72A was analyzed by native mass spectrometry (Native-MS) and size-exclusion chromatography coupled with multi-angle light scattering (SEC-MALS). Both methods showed that, while FAM72A predominantly exists as a monomer in solution, a small fraction forms dimers, with molecular weights and hydrodynamic sizes consistent with these species (Fig. EV4B,C). Hence, FAM72A possesses an intrinsic tendency of dimerization, which can be further induced and strengthened by MKLN1 dimerization within the CTLH E3 complex.

**Figure 3.**
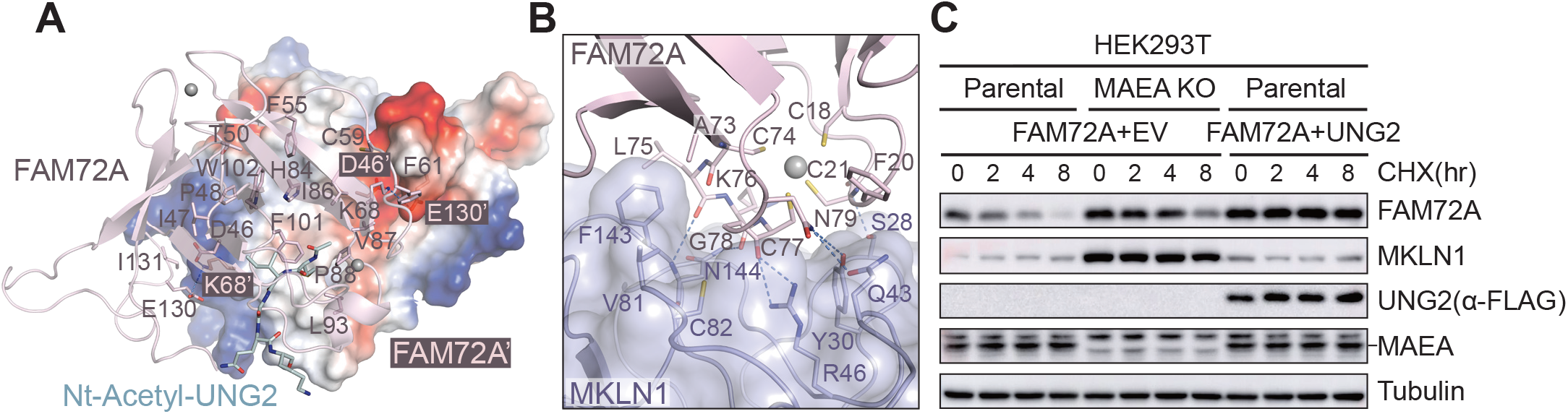
Dimerization and degradation of FAM72A. (A) Top view of the FAM72A dimer interface highlighting a hydrophobic core flanked by electrostatic interactions. The two FAM72A monomers are shown as pink ribbon and surface electrostatic potentials, colored from positive (blue) to negative (red), respectively. For clarity, residues contributing to the hydrophobic core are shown only for one monomer, whereas residues mediating charge interactions (blue dashed lines) are shown for both monomers, with one monomer labeled in a different shade. Nt-Acetyl-UNG2 and zinc ions are shown as teal sticks and gray spheres, respectively. (B) Close-up view of the interface between the discoidin domain of MKLN1 and the zinc finger motif of FAM72A. The discoidin domain is shown as blue surface, and the zinc finger motif is shown as pink ribbon. Selected interface residues are displayed as sticks. Blue dashed lines indicate hydrogen bonds and polar interactions. The zinc ion is shown as a gray sphere, and its four cysteine ligands are shown as sticks. (C) Cycloheximide (CHX) chase assays monitoring the cellular stability of FAM72A. HEK293T parental and MAEA KO cells were transiently co-transfected with the indicated plasmids. 24 h after transfection, cells were treated with CHX for 0-8 h. Whole-cell lysates collected at the indicated time points were analyzed by immunoblotting with the indicated antibodies. EV, empty vector.

As noted earlier, the barrel-shaped discoidin domain of MKLN1 establishes an interface with FAM72A by directly engaging its zinc finger motif and the coordinating loops that stabilize the β-tent fold of FAM72A. Among the two zinc-binding loops, the β6-β7 loop plays the primary role in recognition, with assistance from the β1-β2 loop. In detail, a network of polar interactions involving residues C21, L75, K76, C77, and N79 of FAM72A and residues S28, Y30, Q43, R46, and N144 located on the surface loops of the MKLN1 discoidin domain forms the initial tight association between the two molecules (Figs. 3B and EV3D, 4D). This interaction is further reinforced by hydrophobic contacts involving residues F20, A73, and L75 of FAM72A and residues Q43, V81, C82, and F143 of MKLN1, resulting in a compact yet robust interface (Fig. 3B).

Prior study stated that binding of FAM72A to MKLN1 disrupts MKLN1 tetramerization (Barbulescu *et al*., 2024). The FAM72A-MKLN1 interface revealed in our structure provides the molecular details that mechanistically support this notion. Moreover, MKLN1’s oligomerization has been linked to its degradation by the CTLH E3 ligase in autoregulation (Maitland *et al*, 2024; Yi *et al*., 2024). To test whether physiological expression of FAM72A could protect MKLN1 from degradation, we expressed FAM72A in HEK293T cells at increasing levels and examined steady-state MKLN1 protein abundance. As shown by immunoblotting, FAM72A stabilized MKLN1 in a dose-dependent manner (Fig. EV4E). We next conducted cycloheximide (CHX) chase assays in cells, with or without FAM72A overexpression. Although FAM72A expression increased MKLN1 stability to some extent, we were surprised to find that, the levels of FAM72A declined rapidly over time during the CHX chase assays but were largely preserved in MAEA KO (CTLH E3 catalytic activity-deficient) cells (Figs. 3C and EV4F). This observation implies that FAM72A is regulated and degraded by the CTLH E3 complex, seemingly as a substrate of MKLN1. Considering that, in the case of MKLN1, substrate engagement can stabilize degradable substrate receptor, we next co-expressed UNG2, which binds to FAM72A as a substrate, and observed a dramatic enhancement in FAM72A stability (Fig. 3C). These findings combinedly suggest that, by partnering with MKLN1, FAM72A functions as a dimeric substrate receptor of the CTLH E3 ligase and undergoes stability regulation through CTLH E3-mediated degradation.

### N-terminally acetylated consensus motif as a degron recognized by the CTLH^MKLN1-FAM72A^ E3 ligase

In light of all FAM72A’s features described above, along with its substrate-recognition hinged on a short linear peptide captured in our structural studies, we proposed that FAM72A may represent a CTLH E3 substrate receptor dedicated to degron recognition. To test this idea, we sought to determine the consensus sequence of FAM72A-bound peptide in the first place. Given the strict requirement of the Nt-acetylated methionine at position 1 of the UNG2 peptide for FAM72A recognition, we employed ITC experiments to quantify the binding affinities between FAM72A and a panel of synthesized Nt-acetylated peptides carrying single-site mutations at positions 2−4. Compared with the WT peptide, for the residue I2 which is accommodated in a fitted hydrophobic cavity, substitutions to its hydrophobic analogs L and V, as well as the small uncharged residues like A and T, were generally tolerated and didn’t show drastic influence on the recognition (Fig. 4A). In contrast, replacement with bulky hydrophobic residues such as F or M, or with charged residues, substantially impaired or abolished the binding. As to the residue G3 with a bottleneck-ish spatial constraint in the binding-pocket, substitution with any residue containing a side chain completely eliminated the interaction (Fig. 4B), rendering G as the uniquely required residue at this position. The fourth position was conceivably flexible, as none of the substitutions produced detectable effects on peptide binding (Fig. 4B).

**Figure 4.**
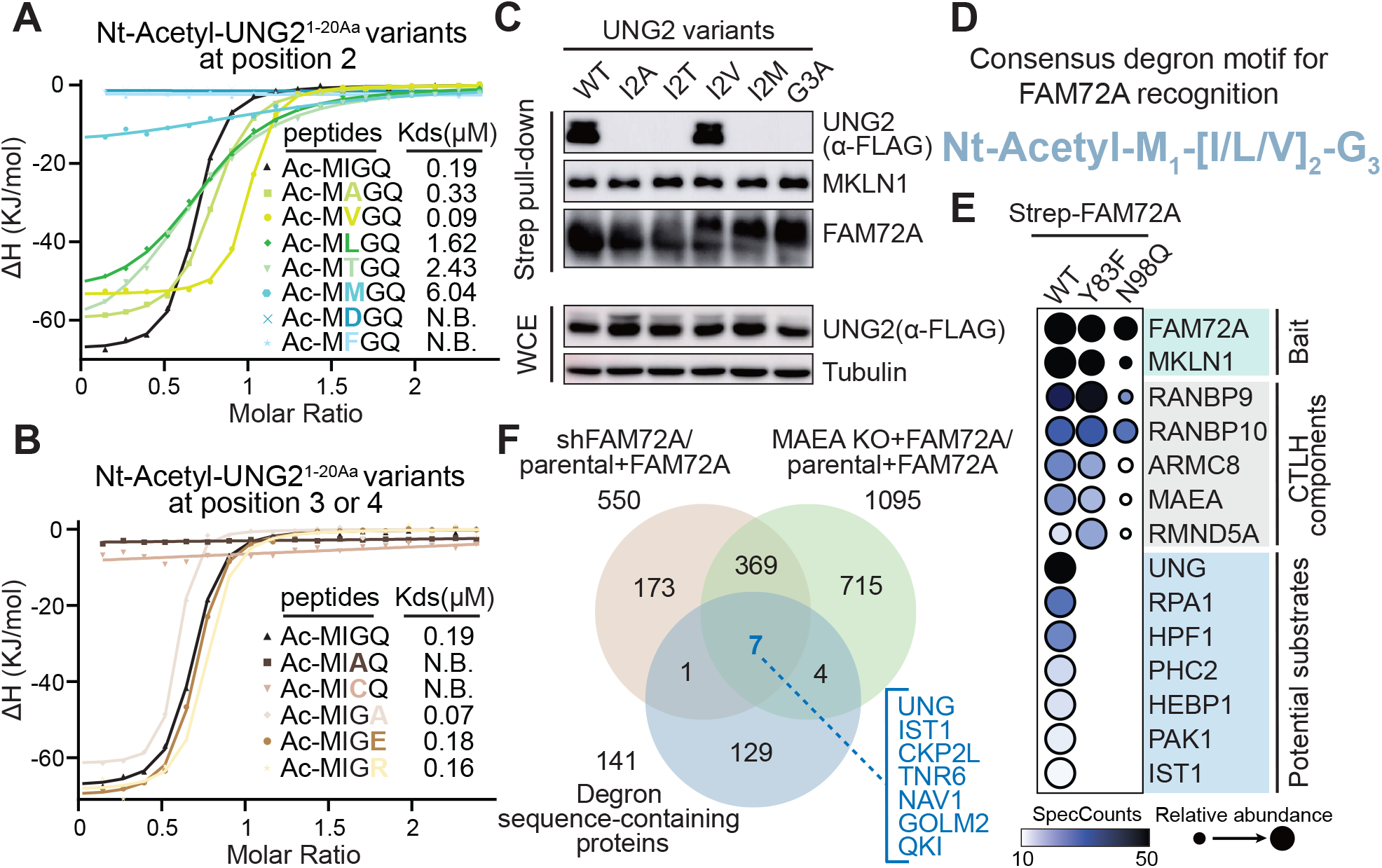
Identification of the consensus degron motif and proteomic screening for potential substrates. (A, B) ITC binding curves for FAM72A titrated with Nt-Acetyl-UNG2 peptides (1-20Aa) containing substitutions at position 2-4. The corresponding peptide sequences and binding affinities are indicated. Substituted residues are highlighted in bold and colored to match the corresponding binding curves. N.B., no detectable binding. (C) Strep pull-down assays assessing the cellular binding of WT UNG2 and its position 2 and position 3 variants to immobilized FAM72A. Whole-cell lysates were used as prey in the pull-down assays. Bound proteins were analyzed by immunoblotting with the indicated antibodies. (D) Consensus N-terminally acetylated degron motif recognized by FAM72A. The motif is shown in blue. Amino acid positions within the motif are indicated as subscripts. Square bracket (“[]”) at position 2 denote that any residue listed within the brackets is tolerated at this position. (E) Dot plot generated using ProHits-viz visualizing the spectral counts of the CTLH complex components and potential substrates captured in AP-MS using WT FAM72A and its UNG2-binding-deficient variants Y83F and N98Q as baits. Dot color and size represent spectral count and relative abundance, respectively. Darker shading indicates higher spectral counts (values >50 are shown in black, while values <10 are shown in white). Dot size reflects the relative abundance of each protein across the three datasets. The bait, CTLH complex components, and potential substrates are grouped and labeled accordingly. (F) Venn diagrams (Heberle *et al*, 2015) showing the overlap of proteins whose stability is regulated by the CTLH^MKLN1-FAM72A^ E3 ligase. Using a *P* value cutoff of 0.05 and a 1.5-fold change threshold, 376 proteins were identified as upregulated in both FAM72A KD cells and MAEA KO cells compared to parental HEK293T cells. Integration of these datasets with the pool of human proteins containing the consensus degron motif identified seven candidate substrates, which are highlighted in sky blue.

It has been documented that, in human cells, the N-terminal initiator methionine is frequently removed co-translationally when followed by a small uncharged residue, giving rise to a processed mature protein lacking the Nt-acetylated methionine at position 1 (Xiao *et al*, 2010). To figure out whether the Nt-acetylated sequences tested in our binding assays are present and recognized by FAM72A in a cellular context, we conducted affinity pull-down experiments using immobilized FAM72A as bait and lysates from HEK293T cells expressing C-tagged UNG2 variants harboring the corresponding mutations analyzed in the peptide-binding studies. With the I2M and G3A mutants as negative controls, we found that only the I2V variant retained interaction with FAM72A at a level comparable to that of the WT protein, whereas neither the I2A nor the I2T mutant displayed any FAM72A-binding activity, indicating that residues A and T are unlikely eligible at position 2 despite being tolerated *in vitro* (Fig. 4C). As such, we biochemically characterized that, instead of relying solely on the Nt-acetylated initiator methionine, the peptide dictating FAM72A recognition features a more stringent sequence context defined by a consensus degron motif of Nt-AcM_1_-[ILV]_2_-G_3_ (Fig. 4D).

To identify potential degron-containing substrates within the human proteome, we employed two complementary proteomics-based screening strategies centered on protein-protein interactions and intracellular protein stability. First, we produced FAM72A-MKLN1 complexes as baits using FAM72A in forms of WT and two substrate-binding-deficient variants identified in our studies, Y83F and N98Q. We next conducted affinity purification followed by mass spectrometry (AP-MS) to examine the FAM72A interactome in the context of the CTLH E3.

Among the 57 FAM72A-interacting proteins identified in the WT sample (spectral counts >10), seven were not detected in either substrate-deficient spectra, suggesting that they may represent candidate substrates (Fig. 4E; Dataset EV2). Notably, except for the well-established substrate UNG2 and its associated RPA1, three of these seven candidates, including HPF1, HEBP1, and IST1, possess explicit degron sequence at their N-termini, manifesting their identities as substrates recognized by FAM72A and its associated CTLH E3 ligase.

Meanwhile, we generated two HEK293T cell lines: an MAEA KO line and a FAM72A KD (shFAM72A) line. FAM72A was subsequently overexpressed in both the parental and MAEA KO cell lines. These cells, together with the FAM72A KD cells, were subjected to quantitative proteomic studies to assess the intracellular protein abundance. Direct data-independent acquisition (DIA) analysis was performed to compare parental cells expressing FAM72A with both FAM72A KD cells and MAEA KO cells expressing FAM72A. Out of the 8202 proteins detected in both comparisons, we identified 550 and 1095 as FAM72A-responsive and MAEA-regulated proteins (p < 0.05 and a fold-change cutoff >1.5), respectively (Fig. 4F; Dataset EV3). By integrating the results from two comparisons with a ScanProsite (Sigrist *et al*, 2013) search for degron-containing proteins in the human proteome (141 hits), we found seven proteins at the intersection as candidate substrates (Fig. 4F). Three candidates were further eliminated because they were transmembrane or secreted proteins or were likely too large to fit within the ring-shaped CTLH E3 complex. This filtering process yielded four proteins, UNG, IST1, QKI, and CKAP2L, as high-confidence substrates. Collectively, the combined proteomic screening approaches identified five candidate substrates for the CTLH^MKLN1-FAM72A^ E3 ligase, including HPF1, HEBP1, IST1, QKI, and CKAP2L.

### Revelation of neo-substrates specific to the AcM-[ILV]-G/N-degron pathway

To validate these candidate substrates in cell-based assays, we utilized the three HEK293T cell lines generated for the quantitative proteomic studies and examined the steady-state protein abundance of the candidate substrates, with UNG2 as a positive control. As opposed to WT cells expressing FAM72A, all candidates except CKAP2L got stabilized and accumulated in both MAEA KO cells expressing FAM72A and FAM72A KD cells, similar to the behavior of UNG2, thereby excluding CKAP2L as a bona fide substrate (Fig. 5A). Meanwhile, to assess substrate recognition, co-IP experiments were performed in cells transfected with either FAM72A WT, or its substrate-deficient variants Y83F and N98Q. To minimize the impact of FAM72A and its variants on substrate abundance and to stabilize the FAM72A-substrate interactions, MAEA KO cells were employed for a clearer assessment. Immunoblot analysis showed that, all four candidates interacted strongly with FAM72A WT, whereas faint to no binding was detected with either substrate-deficient variant (Fig. 5B). This binding pattern mirrored that of UNG2, further evidencing that these proteins are authentic substrates.

**Figure 5.**
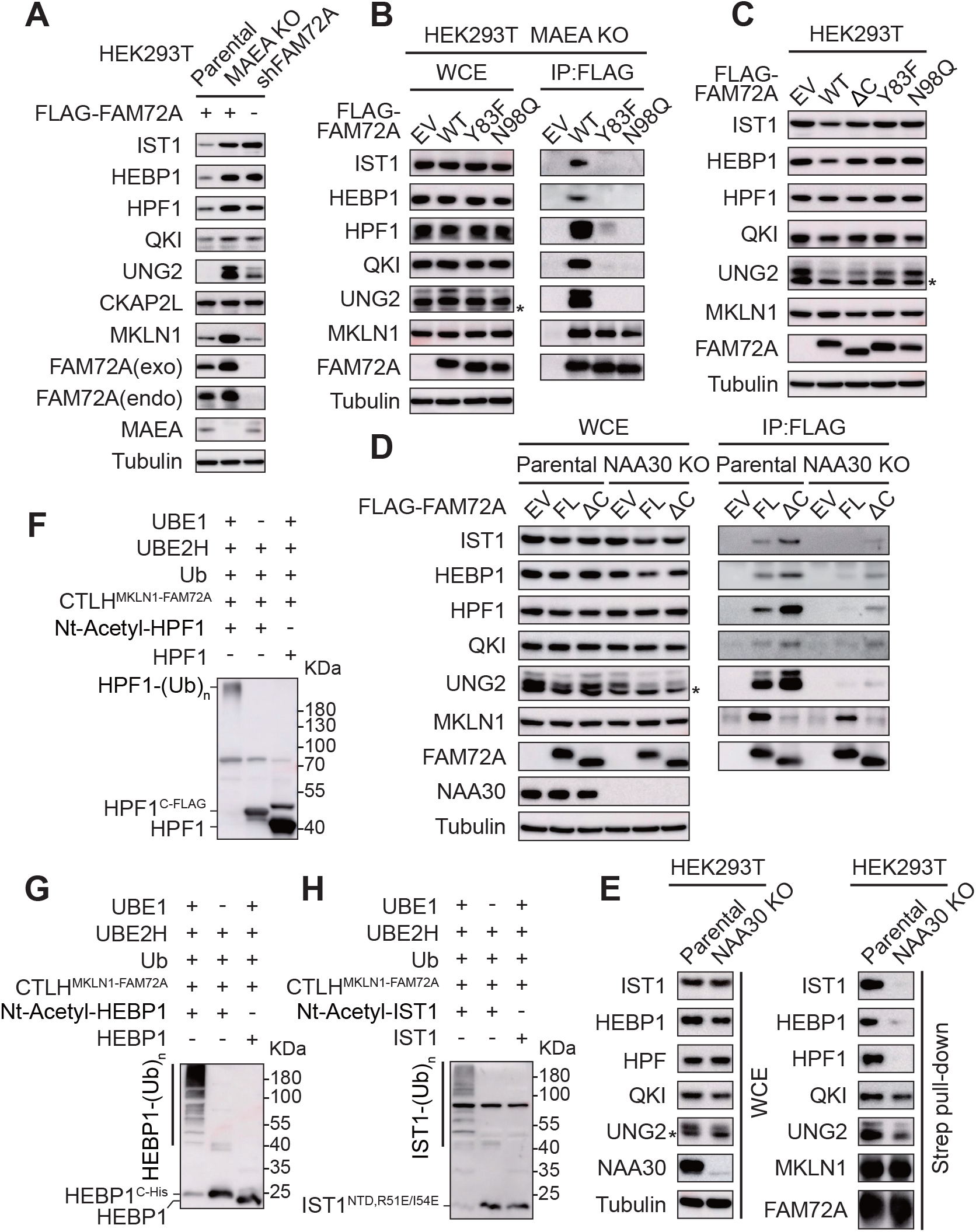
Cellular validation of neo-substrates for the AcM-[ILV]-G/N-degron pathway. (A) Assessment of the steady-state protein abundance of candidate substrates in HEK293T parental, MAEA KO, and FAM72A KD cells. An N-terminally FLAG-tagged FAM72A construct was transiently transfected in the parental and MAEA KO cells. Whole-cell lysates were analyzed by immunoblotting with the indicated antibodies. (B) Cell-based co-IP assays assessing the recognition of candidate substrates by FAM72A and its substrate-binding-deficient variants. HEK293T MAEA KO cells were transiently transfected with the indicated FLAG-tagged FAM72A plasmids. Whole-cell extracts (WCE) were subjected to immunoprecipitation with anti-FLAG beads, followed by immunoblotting with the indicated antibodies. Asterisks denote UNG1 proteins. EV, empty vector. (C) Effect of FAM72A loss-of-function variants on the steady-state protein abundance of candidate substrates in cells. HEK293T cells were transiently transfected with the indicated FLAG-tagged FAM72A plasmids. The ΔC variant of FAM72A comprises the first 144 amino acids. Asterisks denote UNG1 proteins. EV, empty vector. (D) Cell-based co-IP assays assessing the impact of N-terminal acetylation on candidate substrate recognition by FAM72A and its variant. HEK293T parental and NAA30 KO cells were transiently transfected with FLAG-tagged FAM72A constructs in either full-length or ΔC forms. Whole-cell extracts (WCE) were subjected to immunoprecipitation with anti-FLAG beads, followed by immunoblotting with the indicated antibodies. Asterisks denote UNG1 proteins. EV, empty vector. (E) Strep pull-down assays assessing the binding of endogenous candidate substrates to the immobilized FAM72A-MKLN1 complex in HEK293T parental and NAA30 KO cells. Whole-cell lysates were used as prey in the pull-down assays. Bound proteins were analyzed by immunoblotting with the indicated antibodies. Asterisks denote UNG1 proteins. (F-H) *In vitro* ubiquitination of HPF1, HEBP1 and IST1 with or without N-terminal acetylation by the CTLH^MKLN1-FAM72A^ E3 complex. To improve the solubility of IST1, the C-terminal flexible region was removed and two site mutations were introduced into the construct (Bajorek *et al*., 2009). Ubiquitination was detected by immunoblotting with substrate-specific antibodies.

We next evaluated the levels of the candidate substrates in the presence of FAM72A loss-of-function variants. In addition to the substrate-deficient mutants Y83F and N98Q, we included a ΔC variant that has substantially impaired engagement with MKLN1 (Barbulescu *et al*., 2024) and thereby cannot be effectively incorporated into the CLTH E3 complex for substrate ubiquitination and degradation. In line with the behavior of UNG2, while all four candidates exhibited their lowest steady-state levels in cells expressing FAM72A WT, expression of the three functionally flawed FAM72A variants led to stabilization of all candidates, although to varying degrees (Fig. 5C). These results link FAM72A functional deficiency to increased stability of candidates, demonstrating that these proteins are bona fide substrates specific to FAM72A.

Remarkably, MS/MS analysis revealed that three potential substrates identified through the AP-MS approach, HPF1, HEBP1, and IST1, along with UNG2, were meticulously N-terminally acetylated at M1 (Fig. EV5), urging us to investigate whether N-terminal acetylation influences the recognition of all candidates by FAM72A in cells. To address this question, we performed cell-based co-IP assays to evaluate the interactions between candidate substrates, as well as UNG2, and overexpressed FAM72A WT and the degradation-deficient ΔC variant in both WT and NAA30 KO cells. The NAA30 KO cells lack the catalytic subunit of the cognate Nt-acetyltransferase C (NatC) complex, which specifically modifies degron-containing nascent proteins (McTiernan *et al*, 2025), and are therefore deficient in Nt-acetylation. Although input analyses showed moderately decreased levels of UNG2 and HEBP1 in NAA30 KO cells, likely due to degradation through an alternative UBR-mediated pathway (Kim *et al*, 2014; Varland *et al*, 2023), the co-IP results were clear enough (Fig. 5D): In WT cells, all candidates and UNG2 were efficiently recognized by both FAM72A WT and the ΔC variant, with the latter showing stabilized binding. By contrast, loss of Nt-acetylation dramatically disabled substrate recognition.

Complementary *in vitro* pull-down assays were performed using purified FAM72A-MKLN1 complexes as baits and lysates from both WT and NAA30 KO cells. Conforming to the cell-based co-IP results, and with even greater clarity, all four candidates, similar to UNG2, substantially lost interactions with FAM72A when their Nt-acetylation were blocked (Fig. 5E). Additionally, *in vitro* ubiquitination assays demonstrated that, for HPF1, HEBP1, and IST1, only their N-terminally acetylated forms were efficiently polyubiquitinated by the reconstituted E3 complex, whereas the corresponding unmodified versions remained poorly polyubiquitinated (Fig. 5F-H), further proving that targeting of these candidates by FAM72A is strictly dependent on their Nt-acetylation. Altogether, our findings revealed and validated a set of neo-substrates, including HPF1, HEBP1, IST1, and QKI, that are specifically recognized and targeted by the AcM-[ILV]-G/N-degron pathway defined in this study.

### Protein degradation mediated by the AcM-[ILV]-G/N-degron pathway is conditional

The known Ac/N-degron pathways have long been considered conditional and implicated in both protein quality control and the maintenance of protein complex stoichiometry (Park *et al*, 2015; Shemorry *et al*., 2013). To investigate whether protein degradation mediated by the AcM-[ILV]-G/N-degron pathway is regulated, we focused on UNG2, which exerts base excision DNA repair during active DNA replication at replication forks by directly binding to PCNA through its flexible N-terminal region (Otterlei *et al*, 1999). Primary sequence analysis revealed that the canonical PCNA binding motif (PIP-box) spans residues 4-11 of UNG2, immediately adjacent to the AcM-[ILV]-G/N-degron motif (Fig. 6A). This close proximity prompted us to wonder whether engagement of UNG2 with PCNA during its function could shield the degron from recognition? To address this question, we generated a model of UNG2-PCNA complex by AlphaFold 3 (Abramson *et al*, 2024) in alignment with our structure of the UNG2-FAM72A recognition (Fig. 6B). Examination of the binding modes of UNG2 N-terminal region in both complexes revealed that Q4 of UNG2, the first conserved residue in the PIP-box motif, inserts into a so-called Q-pocket at the UNG2-PCNA interface. By contrast, in the UNG2-FAM72A complex, the side chain of Q4 leans against the opening of the FAM72A substrate binding pocket. This apparent steric incompatibility prevents residue Q4 from engaging PCNA and FAM72A simultaneously, rendering the interactions of UNG2 with PCNA and FAM72A mutually exclusive. We next quantified the binding affinity of the N-terminally acetylated UNG2 peptide for PCNA (∼3.3μM) (Fig. 6C) and compared it with the previously determined affinity of the same peptide for FAM72A (∼190nM). The UNG2 peptide bound to PCNA more than 15-fold weaker than to FAM72A, suggesting that FAM72A may be able to displace UNG2 from PCNA.

**Figure 6.**
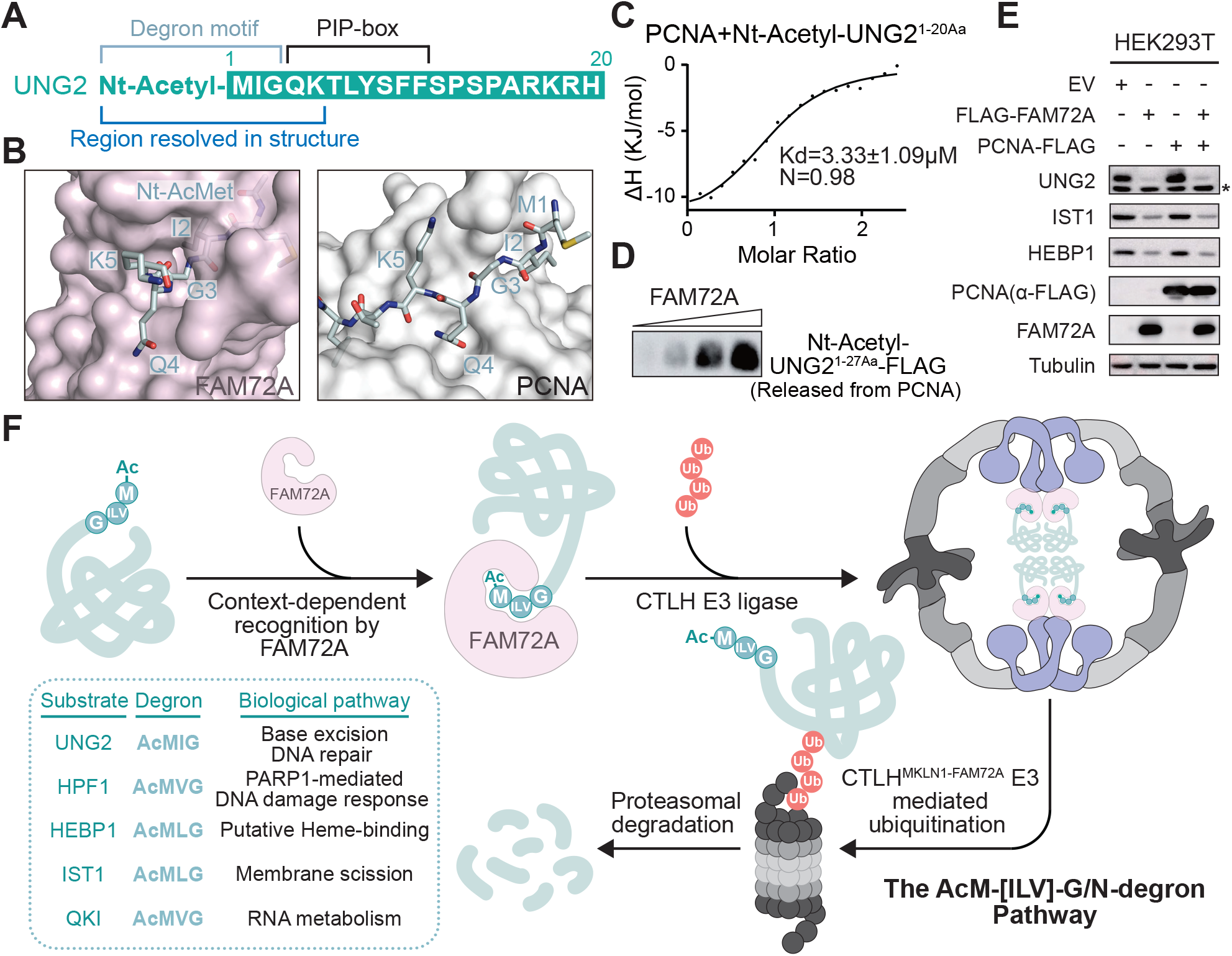
Context-dependent UNG2 degradation and a proposed working model of the AcM-[ILV]-G/N-degron pathway. (A) Primary sequence annotation of the N-terminal region of UNG2. The sequence is shown in teal, with the degron motif, PCNA binding site, and structurally resolved fragment indicated. (B) Side-by-side close-up views of the N-terminal peptide of UNG2 bound to FAM72A (left panel) and PCNA (right panel). The UNG2-PCNA complex model was generated using AlphaFold 3. FAM72A and PCNA are shown as pink and gray surfaces, respectively, and UNG2 peptides are shown as teal sticks with selected residues labeled. (C) ITC binding curve for PCNA titrated with the N-terminally acetylated UNG2 peptide (1-20Aa). The binding affinity is indicated. (D) *In vitro* competition assays demonstrating the ability of FAM72A to disrupt a preformed PCNA-UNG2 complex. An N-terminally acetylated UNG2 peptide (1-27Aa) fused to a C-terminal FLAG tag was synthesized and used in the assay. Dissociation of the peptide from the PCNA-UNG2 complex was monitored in the flowthrough by immunoblotting with a FLAG-specific antibody. (E) Assessment of the steady-state protein abundance of endogenous UNG2 in HEK293T cells. Cells were transiently transfected with the indicated plasmids, and whole-cell lysates were analyzed by immunoblotting with the indicated antibodies. IST1 and HEBP1 were included as controls. Asterisks denote UNG1 proteins. EV, empty vector. (F) Summarized working model of the AcM-[ILV]-G/N-degron pathway defined in this study. All proteins are shown as cartoon representations using the same color scheme as described above. Identified substrates, their corresponding degron motifs, and the biological pathways in which they function are listed.

To test this possibility, we assembled and immobilized the UNG2-PCNA complex on affinity column using purified PCNA and a synthesized Nt-acetylated UNG2 peptide. The complex was then incubated with FAM72A at increasing levels, and the release of the UNG2 peptide into the flow-through was monitored. Immunoblot analysis showed that FAM72A displaced UNG2 from the pre-formed UNG2-PCNA complex in a dose-dependent manner (Fig. 6D). These results were fortified in cell-based assays. Despite the moderate stabilization of UNG2 by PCNA overexpression in HEK293T cells, ectopic expression of FAM72A markedly reduced UNG2 abundance, demonstrating an enhanced activity of the FAM72A-dependent UNG2 degradation (Fig. 6E). In this way, UNG2 degradation mediated by the AcM-[ILV]-G/N-degron pathway is predominantly regulated by the abundance of the substrate receptor FAM72A. When present at elevated levels, FAM72A is competent for dislodging UNG2 from the UNG2-PCNA complex, revealing a role that extends beyond the protein quality control function coupled with the conventional Ac/N-degron pathways.

## Discussion

Degrons act as regulatory elements that couple cellular state to proteome remodeling and determine the selectivity of protein degradation by the UPS. Despite significant advances in degron biology over the past decade (Zhang *et al*., 2025), well-characterized degron-recognin pairs and their recognition and regulatory mechanisms remain to be defined. In this study, we establish a distinct AcM-[ILV]-G/N-degron pathway by identifying a bona fide Ac/N-degron with a consensus Nt-AcM_1_-[ILV]_2_-G_3_ motif, deciphering the mechanism underlying its remarkable recognition specificity, and revealing a cohort of neo-substrates distinctively regulated by this pathway (Fig. 6F). We were particularly impressed by the rigorous selectivity of degron recognition observed in our structural and biochemical analyses. This selectivity went beyond the Nt-acetylated M1 to span the first three N-terminal residues, defining a consensus motif. We realize that such unusual specificity of this degron-recognition system is rooted in a multilayered selection process established during co-translational N-terminal processing of nascent proteins (McTiernan *et al*., 2025) and following substrate receptor recognition. N-terminal methionine excision first removes the initiator methionine followed by a small uncharged residue, so that proteins bearing this sequence pattern are physiologically excluded from recognition. N-terminal acetylation as the second step confers selectivity for an Nt-acetylated methionine at position 1. At the receptor level, FAM72A further sharpens substrate selectivity through a tailored hydrophobic cavity that eliminate large or polar residues at position 2 and a spatially narrowed bottleneck that imposes the most stringent requirement at position 3. Together, these sequential filters, from combined efforts of N-terminal processing and FAM72A, generate the exceptionally strict consensus motif targeted by this degron pathway.

Based on the sequence motif, we initially pinned down 141 degron-containing proteins in the human proteome by database searching. Subsequent proteomic analyses and cell-based validation further narrowed the neo-substrates down to four, comprising a surprisingly small fraction of all degron-containing proteins. What prevents the vast majority of degron-containing proteins from being recognized as substrates? Our structural analyses demonstrated that, in the context of the CTLH E3, FAM72A forms a dimer which is induced and stabilized through MKLN1 dimerization. Within the ring-shaped CTLH E3 assembly, dimerization of the substrate receptor leads to the probability that substrates are recruited and engaged in an oligomeric state, which coincides with the predisposition of the CTLH E3 complex for accommodating oligomeric substrates and with previous observations that the FAM72A-bound UNG2 adopts a dimeric configuration (Barbulescu *et al*., 2024; Gottemukkala *et al*., 2024; Sherpa *et al*., 2021).

Intriguingly, among the four identified neo-substrates, IST1 and QKI have been structurally characterized as an oligomer and a dimer, respectively (Nguyen *et al*, 2020; Teplova *et al*, 2013). By contrast, with no direct evidence currently supporting oligomerization of HPF1 or HEBP1 in any known context, whether they bind FAM72A as oligomeric states warrants further investigation. Nevertheless, the dimerization of FAM72A as a substrate receptor along with the oligomeric/dimeric nature of the identified substrates prompts us to propose that, the propensity to oligomerize or the property of being amenable to FAM72A-induced oligomerization within the hollow center of the CTLH E3 assembly enforces an additional layer of three-dimensional substrate selection beyond the linear consensus degron sequence. Such structural requirements may ultimately determine the authentic substrates of this degron pathway.

As observed throughout the study, the AcM-[ILV]-G/N-degron pathway is mediated at multiple levels, underlining its highly regulated cellular activity and exemplifying the context-dependent conditionality of degron system. At first, the pathway strictly relies on co-translational N-terminal acetylation of the initiator methionine in nascent proteins. Loss of this specific modification in the identified substrates has been demonstrated to abolish substrate recognition and consequently deactivate their ubiquitination. Second, the abundance of substrate receptor FAM72A is regulated to dictate the activity of this pathway. We found that FAM72A itself undergoes stability control via CTLH E3-mediated degradation, revealing a mechanism that mirrors the previously reported autoregulation of the CTLH/GID E3 ligase, in which its own substrate receptor MKLN1 and yGid4 are targeted for degradation (Maitland *et al*, 2019; Menssen *et al*, 2018). Third, and most importantly, the pathway depends on the functional context of its substrates. For example, degradation of UNG2 has been revealed to be driven by its FAM72A-mediated displacement from PCNA, despite the marginal protection of its degron conferred by PCNA binding. IST1 (Increased Sodium Tolerance 1), a critical component of the ESCRT-III (Endosomal Sorting Complex Required for Transport-III) complex, assembles with another ESCRT-III protein, CHMP1B, to form hetero-oligomeric CHMP1B-IST1 filaments that promote membrane tube constriction and membrane scission (Cada *et al*, 2022; Nguyen *et al*., 2020). Unlike other canonical ESCRT-III proteins, which can be maintained in a monomeric, autoinhibited closed conformation before activation, IST1 adopts a closed conformation across both its monomeric and active polymeric states (Bajorek *et al*, 2009; McCullough *et al*, 2015; Nguyen *et al*., 2020). We propose that, to restrict the level of potentially polymerization-competent monomeric IST1, the degron pathway may be employed to strategically target the monomeric form while sparing IST1 incorporated in higher-order assemblies. This model is consistent with the likely inaccessibility of the degron observed in the structure of the CHMP1B-IST1 complex (Fig. EV6A) (McCullough *et al*., 2015; Nguyen *et al*., 2020), as well as with the stabilization of IST1 upon co-expression of CHMP1B in the CHX chase assays in cells (Fig. EV6B). Nevertheless, the precise roles of this degron pathway in mediating the functions of these substrates remains to be elucidated.

Lately, the CTLH E3 ligase has emerged as an attractive platform for therapeutic targeted protein degradation (TPD). The modular substrate receptor architecture and the distinct expression patterns of various CTLH E3s offer the potential for more selective and tunable degradation strategies, which has been exemplified by the ligand discovery and the development of TPD strategies focused on the canonical substrate receptor GID4 (Chana *et al*, 2026; Li *et al*, 2025), as well as by recent molecular glue design that harnesses the ligand-binding pocket of the YPEL5, a reported modulator of another substrate receptor WDR26 (Gottemukkala *et al*., 2024; Zhuang *et al*, 2026). Regarding the AcM-[ILV]-G/N-degron pathway, its specific recognin FAM72A, also functioning as a substrate receptor of the CTLH E3, shares structural homology with CRBN-CTD and YPEL5 (Fig. EV3A-C), whose ligand-binding pockets have been successfully exploited for TPD application (Oleinikovas *et al*, 2024; Zhuang *et al*., 2026). By analogy, FAM72A and the generally hydrophobic AcM-[ILV]-G/N-degron it recognizes represent a promising recognition module for the development of novel TPD strategies.

Strikingly, although FAM72A has very restricted physiological expression in most normal tissues, it is frequently overexpressed across a wide range of human malignancies, including breast, colon, and lung cancers (Feng *et al*., 2025; Guo *et al*., 2008). This tumor-associated upregulation of FAM72A potentiates its utilization to achieve highly selective TPD. Furthermore, the structural preference of the FAM72A-degron module within the CTLH E3 assembly for substrates in oligomeric states underscores its potential of TPD applications against diseases in which oligomerization is essential for pathogenic function. In a broader context, our findings not only reshape the current understanding of the conventional Ac/N-degron system, but also open opportunities for developing TPD strategies with intrinsic selectivity embedded in specific E3-degron recognition module.

## Acknowledgments

We thank Wanwan Hu and Hui Shi at Biortus for help on cryo-EM data collection and processing, Mengsi Sun for technical support in ITC experiments, Cookson K. C. Chiu for assistance in LC-MS/MS analysis, and the members of Wang lab for discussion and help. We are grateful to the Core Facility at Shenzhen Bay Laboratory. This work is supported by grants from the National Natural Science Foundation of China (22577079 to H.W.) and the Shenzhen Bay Laboratory Start-up Funds (21320081 to H.W.).

## Author contributions

**Maosheng Yang**: Data curation; Formal analysis; Investigation; Methodology; Resources; Validation; Visualization; Writing—review and editing. **Shuqi Yao**: Data curation; Formal analysis; Investigation; Methodology; Resources; Validation; Visualization; Writing—review and editing. **Zheng Fang**: Investigation; Methodology; Validation. **Wenjie Ren**: Investigation; Methodology; Validation. **Yanyang Yang**: Investigation; Methodology; Validation. **Qizhou Song**: Investigation. **Wen Yang**: Conceptualization; Formal analysis; Methodology; Project administration; Resources; Validation; Visualization; Writing—review and editing. **Hui Wang**: Conceptualization; Formal analysis; Funding acquisition; Methodology; Resources; Supervision; Visualization; Writing—original draft; Writing—review and editing.

## Methods

### Protein expression and purification

Human FAM72A was expressed in *E. coli* Rosetta (DE3) cells as an N-terminal 6×His-GB1 (B1 domain of Protein G) fusion protein in LB medium supplemented with 50 μM zinc sulfate.

Following cell lysis, the fusion protein was purified by nickel affinity chromatography, and the affinity tag was removed by off-column cleavage with tobacco etch virus (TEV) protease. The cleaved protein was further purified by anion exchange chromatography (HiTrap Q HP; Cytiva) followed by size-exclusion chromatography (Superdex 200 Increase; Cytiva). Human UNG2 and PCNA, used as bait proteins in pull-down assays, were expressed in *E. coli* as non-cleavable 6×His fusion proteins and purified by nickel affinity chromatography followed by ion exchange (HiTrap Q HP or HiTrap SP HP as appropriate; Cytiva) and size-exclusion chromatography (Superdex 200 Increase; Cytiva). All the protein mutants and variants were expressed and purified using the same procedures as their corresponding WT proteins.

For preparation of fully phosphorylated UBE2H used in the ubiquitination assays, GB1-tagged UBE2H was co-expressed with the catalytic subunit of CK2 kinase (CSK21) in *E. coli* BL21 (DE3) cells. To generate substrate proteins (UNG2, HEBP1, and IST1^1-194Aa, R51E/I54E^) with an unmodified initiator methionine, an N-terminal 6×His-SUMO tag was introduced and subsequently removed by cleavage with SUMO protease ULP1. Phospho-UBE2H and the substrate proteins were purified using the same procedures described above.

The human CTLH catalytic module (MAEA-RMND5A) and scaffolding module (RANBP9^Δ1-134, Δ468-600^-TWA1) were co-expressed in *E. coli* BL21 (DE3) cells and purified by nickel affinity chromatography followed by anion exchange and size-exclusion chromatography (Superose 6 Increase; Cytiva). To reconstitute the substrate-bound MKLN1-FAM72A complex, MKLN1 was expressed as an N-terminal maltose-binding protein (MBP) fusion protein, while non-tagged substrate proteins were co-expressed with His-tagged FAM72A. Human HPF1 was expressed as a C-terminal FLAG-tagged fusion protein in mammalian cells. HEK293F cells were cultured in OPM-293 CD05 Medium at 37°C in a humidified atmosphere containing 5% CO_2_ with shaking at 100 rpm. Cells were transfected at a density of 1.0×10^6^ cells/mL with plasmid DNA using polyethylenimines (PEI; FUSHENBio) at a final concentration of 1 μg/mL. Following transfection, cells were cultured for an additional 48 h before harvest. Harvested cells were lysed by mild sonication in lysis buffer containing 20 mM Tris-HCl pH 7.5, 150 mM NaCl, supplemented with protease inhibitors (aprotinin, leupeptin, and pepstatin A; 1 μg/mL each). Cell debris and insoluble fraction were removed by centrifugation at 18,000 rpm for 90 min at 4°C. The cleared lysate was incubated with anti-FLAG M2 affinity gel (Sigma-Aldrich) for 2 h, followed by extensive washing with lysis buffer. FLAG-tagged HPF1 was eluted using FLAG peptide at 0.2 mg/mL and subsequently reconstituted with purified FAM72A. Purified MKLN1 and FAM72A-substrate complexes were mixed at a 1:1 molar ratio and further purified by size-exclusion chromatography (Superose 6 Increase; Cytiva). For cryo-EM analysis, the preassembled UNG2-FAM72A-MKLN1 complex was mixed with the RANBP9-TWA1 module at a molar ratio of 1:3 and subjected to a final size-exclusion chromatography (Superose 6 Increase; Cytiva) to separate excess RANBP9-TWA1 complex.

### Affinity pull-down assay

Affinity pull-down assays were performed using purified 6×His-tagged UNG2 WT protein or its variants as bait and purified FAM72A as prey. Approximately 100 μg of bait protein was mixed with ∼200 μg of prey protein and incubated with 100 μL of His-tag purification resin (Roche) at 4°C for 1 h. The resin was then washed extensively with binding buffer to remove unbound proteins. Bound proteins and protein complexes were eluted with binding buffer containing 250 mM imidazole. Eluted samples were resolved by SDS-PAGE and analyzed by Coomassie staining. For Strep-tag pull-down assays, purified Strep-tagged proteins were immobilized on the pre-equilibrated Streptavidin Magnetic Beads (Thermo Scientific) as baits to capture interacting proteins from ∼10 mg of total protein in the indicated precleaned cell lysates by incubation for 2 h at 4℃. After extensive wash with lysis buffer, the beads were analyzed with SDS-PAGE followed by immunoblotting using the indicated antibodies.

### *In vitro* competition assay

Bead-based competition assays were performed to access the ability of FAM72A to displace UNG2 from PCNA. Purified His-tagged PCNA was immobilized on nickel resin and incubated with an excess of N-terminally acetylated UNG2^1-27Aa^ peptide carrying a C-terminal FLAG tag to assemble the PCNA-UNG2 complex. Following extensive washing with binding buffer, the resin was aliquoted into 200 μL (1:1 slurry; ∼20 nmol protein complex) and transferred to a panel of 7.5-mm gravity-flow columns. The immobilized complexes were incubated with 300 µL of purified FAM72A at 0.625, 2.5, 10, or 40 nmol for 1 h at 4℃. The flow-through and a subsequent one-column-volume wash fraction were collected to monitor the dissociation of FLAG-tagged UNG2 from the pre-formed PCNA-UNG2 complex. Samples were analyzed with SDS-PAGE followed by immunoblotting using an anti-FLAG antibody.

### Isothermal titration calorimetry (ITC)

All protein samples used in ITC underwent buffer exchange to 1×PBS via size exclusion chromatography as the last step of purification. All the peptides used in this study were synthesized by GenScript. The lyophilized peptides were dissolved in 1×PBS and desalted using PD MiniTrap G-10 columns (Cytiva). Peptide concentrations were determined by the absorbance at 205 nm using an average absorptivity of 31 mL·mg^-1^·cm^-1^. ITC measurements were performed at 20℃ with a PEAQ-ITC Automated calorimeter (Malvern Panalytical). The typical titration was carried out with 20 µM PCNA or FAM72A WT or its variants in the cell as titrate and 250 µM indicated peptides in the syringe as titrant. Data were analyzed by MicroCal PEAQ-ITC analysis software and fitted to the one set of sites binding model.

### Native mass spectrometry (Native-MS)

Protein samples were buffer-exchanged into 100 mM ammonium acetate prior to analysis. Intact mass spectra were acquired by size-exclusion chromatography-mass spectrometry using an Agilent G6230B TOF mass spectrometer equipped with a dual Agilent Jet Stream electrospray ionization source. The following instrument settings were used in positive ion mode: drying gas temperature, 120 °C; sheath gas temperature, 120 °C; drying gas flow rate, 10 L/min; sheath gas flow rate, 10 L/min; nebulizer pressure, 30 psi; capillary voltage, 5000 V; nozzle voltage, 2000 V; fragmentor voltage, 250 V; and skimmer voltage, 100 V. Spectra were obtained over an *m/z* range of 600-7000 in profile mode and analyzed using Agilent BioConfirm 10.0.

### Size-exclusion chromatography coupled with multi-angle light scattering (SEC-MALS)

SEC-MALS was performed using a system comprising a Shimadzu LC-2025C HPLC for solvent delivery and UV detection, a DAWN multi-angle light scattering detector (Wyatt Technology), and an Optilab differential refractive index detector (Wyatt Technology). A total of 100 μg of purified protein was loaded onto a Superdex 200 Increase 10/300 GL column (Cytiva) and eluted at a flow rate of 0.5 mL/min with buffer containing 50 mM Tris-HCl pH 7.5 and 300 mM NaCl. The refractive index increment (dn/dc) was assumed to be 0.185 mL/g for all protein samples.

Data were collected and analyzed using ASTRA 7.3.2 software. Bovine serum albumin (BSA) was used to calibrate the system.

### Affinity purification mass spectrometry (AP-MS) and data analysis

N-terminally 3×StrepII-tagged FAM72A was expressed and purified using the same procedure described above for GB1-tagged protein. The WT FAM72A-MKLN1 complex and two substrate-binding-deficient variants (FAM72A^Y83F^-MKLN1 and FAM72A^N98Q^-MKLN1) were assembled and purified based on the N-terminally 3×StrepII-tagged FAM72A. Purified FAM72A or the corresponding protein complexes were immobilized on pre-equilibrated Streptavidin Magnetic Beads (Thermo Scientific) as baits. HEK293T cell lysates were precleaned by streptavidin magnetic beads to remove biotin-binding and non-specifically binding proteins. Approximately 20 µL of bait-bound beads were incubated with ∼10 mg of total protein from the pre-treated lysate at 4℃ for 2 h. After extensive wash with lysis buffer, the beads were applied to SDS-PAGE for analysis. To prepare samples for mass spectrometry, the SDS-PAGE gel covering all protein bands of interest was excised. Subsequent in-gel tryptic digestion, peptide extraction, LC-MS/MS data acquisition, and preliminary analysis were performed by Omicsolution Co., Ltd.

Raw proteomic data were searched against the UniProt human proteome (20,656 entries; 2025 release). Peptide and protein identifications were filtered to a 1% false discovery rate (FDR), and proteins identified by at least one unique peptide were retained for further analysis. To identify candidate substrates of the CTLH^MKLN1-FAM72A^ E3 ligase, proteins were required to have no spectral counts in either of the two substrate-binding-deficient variants and more than 10 spectral counts in the WT complex. Spectral counts for known CTLH E3 components and high-confidence candidate substrates were visualized as dot plots generated by ProHits-viz (Knight *et al*, 2017).

### N-terminal modification analysis

Protein samples were separated by SDS-PAGE, and gel regions corresponding to the bands of interest were excised for subsequent MS/MS analysis. The acquired MS/MS spectra were searched against the UNG2 protein sequence and a contaminant database (UniProt human proteome described above). Carbamidomethylation of cysteine was defined as a fixed modification, whereas oxidation of methionine, deamidation of asparagine and glutamine, and protein N-terminal acetylation were set as variable modifications. MS/MS fragmentation spectra were located and interpreted using Qual Browser within Xcalibur software (Thermo Fisher Scientific). Theoretical fragment ion masses were generated using MS-Product in the UCSF Protein Prospector package. A summary of the theoretical and experimentally observed fragment ion masses, together with the corresponding mass errors, is provided in Dataset EV1.

### *In vitro* ubiquitination assay

Ubiquitination reactions were performed using 0.5 µM UBE1 (Biortus), 2 µM phospho-UBE2H, 20 µM ubiquitin, 0.5 µM of the CTLH central catalytic and scaffolding module (MAEA-RMND5A-RANBP9-TWA1), and 0.5 µM of the CTLH substrate recognition module-substrate complex (AcUNG2-FAM72A-MKLN1) in reaction buffer containing 20 mM Tris-HCl pH 7.5, 150 mM NaCl, 5 mM ATP, and 10 mM MgCl_2_. Control reactions were performed either in the absence of UBE1 or using a non-N-terminally acetylated form of UNG2. Reaction mixtures were incubated at 30 °C for 1.5 h and quenched with SDS sample loading buffer. Ubiquitination assays using HEBP1, IST1^1-194Aa, R51E/I54E^, and HPF1 as substrates were performed under identical conditions. Ubiquitin conjugates were detected by immunoblotting with the indicated antibodies.

### Cryo-EM sample preparation and data collection

UltrAuFoil 300 mesh R1.2/1.3 grids (Quantifoil Micro Tools GmbH) were glow-discharged at 15 mA for 100 s using a PELCO easiGlow glow discharge system. 3 μL of the purified UNG2-FAM72A-MKLN1-RANBP9-TWA1 complex (2.0 mg/mL) was applied to a freshly glow-discharged grid. The grid was subsequently blotted for 3-5 s (blot force 0, 4 °C, and 100% relative humidity) and plunge-frozen in liquid ethane using a Vitrobot Mark IV system (Thermo Fisher Scientific). Frozen grids were stored in liquid nitrogen until cryo-EM data collection.

Cryo-EM data were collected on a Titan Krios G4 transmission electron microscope operated at 300 kV and equipped with a Gatan K3 direct electron detector operating in super-resolution bin 2 mode. An energy filter with a slit width of 20 eV was used. Movies were recorded in TIF format using EPU (Thermo Fisher Scientific) at a nominal magnification of 105 K, resulting in a physical pixel size of 0.821 Å. Images were recorded over a defocus range of -1.0 to -2.0 μm with a total electron exposure of 60 e^-^/Å^2^.

### Image processing and 3D reconstruction

Cryo-EM image processing was performed using CryoSPARC (Punjani *et al*, 2017) and RELION (Zivanov *et al*, 2018). A total of 5,005 micrographs were retained after motion correction, CTF estimation, and manual inspection. Particles (n=7,403,304) were automatically picked using the Blob Picker, Template Picker, and Topaz (Bepler *et al*, 2019) algorithms implemented in CryoSPARC. Following data cleaning by 2D classification, 2,521,495 particles were subjected to two-round *ab-initio* reconstruction and heterogeneous refinement. A subset of 213,248 particles corresponding to the best-resolved classes was selected for 3D classification and non-uniform refinement (Punjani *et al*, 2020), resulting in a reconstruction at 3.19 Å resolution. The selected particles were subsequently processed in RELION through 3D auto-refinement, Bayesian polishing, and post-processing, yielding a reconstruction at 2.89 Å resolution. Lastly, C2 symmetry, non-uniform refinement, and local refinement were applied to further improve the map quality. The final reconstruction reached an overall resolution of 2.81 Å, as determined by the gold-standard FSC (Fourier shell correlation) using the 0.143 criterion. Local resolution was estimated in the CryoSPARC, and the final density map was sharpened using a B-factor of -60.6 Å^2^.

### Model building and refinement

Initial structural models of MKLN1 and FAM72A were generated by AlphaFold 3 (Abramson *et al*., 2024) and rigid-body fitted into the 2.81 Å cryo-EM map using UCSF ChimeraX (Meng *et al*, 2023). The FAM72A-MKLN1 complex was modeled as a C2-symmetric dimer based on the reconstructed cryo-EM map. Two prominent, unassigned densities were observed in FAM72A, each coordinated by the side chains of four cysteine residues, consistent with the geometry of a metal-binding site. These densities were assigned as two zinc ions, supported by the molecular weight determined by native mass spectrometry. For the substrate UNG2, density was only observed for the N-terminal acetylated pentapeptide, which was built into the model. The RANBP9-TWA1 module was not modeled due to the lack of density. The model was iteratively improved through rounds of real-space refinement in PHENIX (Liebschner *et al*, 2019) and manual adjustment in Coot (Emsley *et al*, 2010). The final model exhibited 94.93% of residues in favored regions and 5.07% in allowed regions of the Ramachandran plot. Data collection, model refinement and validation statistics are summarized in Table EV1. All structure figures were rendered in PyMOL or ChimeraX.

### Mammalian cell culture

HEK293T cell lines were purchased from MeilunBio and routinely monitored for *Mycoplasma* contamination using the Universal Mycoplasma Detection Kit (Yeasen Biotechnology). Cells were maintained in Dulbecco’s modified Eagle’s medium (DMEM; Gibco) supplemented with 10% fetal bovine serum (FBS; Sigma-Aldrich) and 1% Penicillin-Streptomycin (Gibco) at 37°C in a humidified incubator with 5% CO_2_.

### CRISPR-Cas9-mediated genome editing

HEK293T knockout (KO) cell lines were generated by CRISPR-Cas9-mediated genome editing. Single-guide RNAs (sgRNAs) targeting Exon 2 of MAEA (CGTTTGTTCAGCGTCTCGTA) (Lampert *et al*, 2018; Maitland *et al*., 2019) and Exon 2 of NAA30 (sgRNA-1: GCCGGGGTACACTCGGGCGAG, sgRNA-2: GACCGCCGACTGCAGCTTAA) were designed and individually cloned into the pSpCas9(BB)-2A-GFP (PX458) vector. HEK293T cells were cultured to approximately 70% confluency and transfected with the indicated plasmids using Lipofectamine 3000. 48 h post-transfection, GFP-positive single cells were isolated by fluorescence-activated cell sorting (FACSAria III, BD) into 96-well plates and grow. Genomic DNA was extracted from individual clones and subjected to genotyping PCR using the following primer pairs: MAEA, F-5’-CATCCAGGGATGTGGAGTGG and R-5’-ATTGCAGCTTGCCTGTCTTAGG; NAA30, F-5’-AGCAGCAGCTCAACGGATTGAT and R-5’-GATCACTGCTGGGTTCACCCTC. PCR products from candidate clones were purified and sequenced to identify clones harboring disruptive mutations. Gene disruption in positive clones was further confirmed by immunoblotting.

### Lentivirus production and stable cell line generation

To generate FAM72A knockdown (KD) HEK293T cell lines, an shRNA targeting FAM72A (CCAGGCAGTTTATGATATTAA), designed using the Sigma Predesigned shRNA tool, was cloned into the pLKO.1 lentiviral vector. Recombinant pLKO.1, psPAX2, and pMD2.G plasmids were co-transfected into HEK293T cells at a ratio of 3:1:1 using polyethylenimine (PEI; FUSHENBIO) according to the manufacturer’s instructions. Lentiviral supernatants were collected 48 h post-transfection and passed through a 0.22 μm sterile filter (Millipore) to remove cellular debris. HEK293T cells were transduced with the filtered lentiviral supernatants in the presence of 10 μg/mL polybrene for 24 h. Following transduction, cells were selected with 2 μg/mL puromycin for one week, and pooled puromycin-resistant cells were validated by immunoblotting.

### Transient expression in mammalian cells

cDNAs encoding FAM72A WT and its variants were cloned into a modified pcDNA3.0 vector containing an N-terminal 3×FLAG tag. cDNAs encoding PCNA, IST1, UNG2 WT, and its mutants were cloned into the pcDNA3.1 vector to generate proteins with a C-terminal 3×FLAG tag. cDNA encoding CHMP1B was cloned into the pcDNA3.1 vector to generate non-tagged protein. HEK293T cells were transiently transfected with the indicated plasmids using polyethylenimine (PEI; FUSHENBIO) according to the manufacturer’s instructions. Cells were harvested 36 h after transfection, and whole-cell lysates were prepared for subsequent analyses.

### Immunoprecipitation and immunoblotting

Indicated cells were harvested and lysed on ice in lysis buffer (Beyotime) supplemented with a protease inhibitor cocktail (Sigma-Aldrich). Cell lysates were clarified by high-speed centrifugation for 30 min at 4°C, and protein concentrations were determined using the Pierce BCA Protein Assay Kit (Thermo Scientific) according to the manufacturer’s instructions. For immunoprecipitation of FLAG-tagged proteins, clarified lysates were incubated with pre-equilibrated anti-FLAG magnetic agarose (Thermo Scientific) for 2 h at 4°C. Beads were subsequently washed extensively with lysis buffer.

Whole-cell lysates and the immunoprecipitated proteins were resolved by SDS-PAGE and transferred onto PVDF membranes (Millipore). Membranes were stained with Ponceau S (Beyotime) to verify protein transfer, blocked with 5% (w/v) non-fat milk in PBST for 1 h at RT, and incubated with the indicated primary antibodies overnight at 4°C. Following incubation with the appropriate horseradish peroxidase-conjugated secondary antibodies, immunoreactive bands were visualized using SuperSignal West Pico PLUS Chemiluminescent Substrate (Thermo Scientific) and imaged with an Amersham ImageQuant 800 system (Cytiva).

### Quantitative proteomics and data analysis

To identify candidate substrates of the CTLH^MKLN1-FAM72A^ E3 ligase, three HEK293T cell lines were analyzed: WT cells overexpressing FAM72A, MAEA KO cells overexpressing FAM72A, and FAM72A KD cells. Total protein was extracted from approximately 10^7^ cells per condition, followed by denaturation, reduction, alkylation, and tryptic digestion. Peptides were desalted using C18 cartridges, and 200 ng of peptide from each sample was analyzed by LC-MS/MS. Peptides were separated on an AUR3-15075 C18 analytical column using a Vanquish Neo UHPLC system (Thermo Fisher Scientific) connected to a timsTOF HT mass spectrometer (Bruker). Peptide elution was executed by a binary buffer system consisting of buffer A (0.1% formic acid) and buffer B (80% acetonitrile with 0.1% formic acid). The gradient was programmed from 2.2% to 44% buffer B over 15 min, followed by an increase to 90% buffer B over 3 min and a 3-min wash at 90% buffer B. The column temperature was maintained at 50°C. MS data were acquired using the diaPASEF (parallel accumulation-serial fragmentation combined with data-independent acquisition) acquisition mode (Meier *et al*, 2020). A total of 82 × 10 Th precursor isolation windows spanning an *m/z* range of 380.8 to 1170.8 were defined, and 3-6 repetitions were applied within a 16-scan diaPASEF scheme to adapt the MS1 cycle time.

DIA raw data were processed and analyzed using Spectronaut 20 (Biognosys AG). Protein identification and quantification were performed against the UniProt human proteome (20,656 entries; 2025 release). Trypsin was specified as the proteolytic enzyme, allowing up to two missed cleavages, and peptides with lengths ranging from 7 to 52 amino acids were considered. Carbamidomethylation of cysteine residues was set as a fixed modification, whereas methionine oxidation and protein N-terminal acetylation were specified as variable modifications. The false discovery rate (FDR) was controlled at 1% at the precursor, peptide, and protein levels. Protein quantification was determined using the Local Normalization strategy and MaxLFQ algorithm.

### Cycloheximide (CHX) chase assay

Cellular protein stability was assessed using a cycloheximide (CHX) chase assay. HEK293T parental and/or MAEA KO cells were transiently transfected with the indicated plasmids for 24 h. Cells were then treated with 100 μg/ml of CHX (Selleck) to inhibit de novo protein synthesis and harvested at the indicated time points post-treatment. Whole-cell lysates were prepared and analyzed by immunoblotting.

### Antibodies

Antibodies used for immunoblotting were anti-UNG (1:3000, Abcam ab245630), anti-IST1 (1:5000, Proteintech 51002-1-AP), anti-HPF1 (1:5000, Proteintech 28456-1-AP), anti-HEBP1 (1:5000, Proteintech 82950-1-RR), anti-QKI (1:5000, Proteintech 13169-1-AP), anti-CKAP2L (1:2500, Proteintech 17143-1-AP), anti-FAM72A (1:2000, Proteintech 33987-1-AP), anti-NAA30 (1:3000, Proteintech 25149-1-AP), anti-MKLN1 (1:1000, Santa Cruz Biotechnology sc-398956), anti-MAEA (1:1000, Abcam ab151304), anti-CHMP1B (1:4000, Proteintech 14639-1-AP), anti-β-Tubulin (1:1000, Cell Signaling Technology #15115), anti-DYKDDDDK Tag (1:1000, Cell Signaling Technology #14793), anti-rabbit IgG, HRP-linked antibody (1:3000, Cell Signaling Technology #7074), and anti-mouse IgG, HRP-linked antibody (1:3000, Cell Signaling Technology #7076).

### Quantification and statistical analysis

Protein concentrations were determined using the Bradford Protein Assay protocol and Bio-rad Protein Assay Dye with a NanoDrop One^C^ spectrophotometer (Thermo Fisher Scientific).

Statistical analysis of the proteomics data was performed on the basis of three biological replicates. Statistical significance was assessed using a two-tailed unpaired Student’s t-test.

## Data availability

The cryo-EM density map of the human UNG2-FAM72A-MKLN1 complex has been deposited in the Electron Microscopy Data Bank (EMDB) with the accession code: EMD-82509. The corresponding atomic coordinates have been deposited in the Protein Data Bank (PDB) with the accession code PDB: 44CV.

## Expanded View Figure Legends

**Figure EV1.**
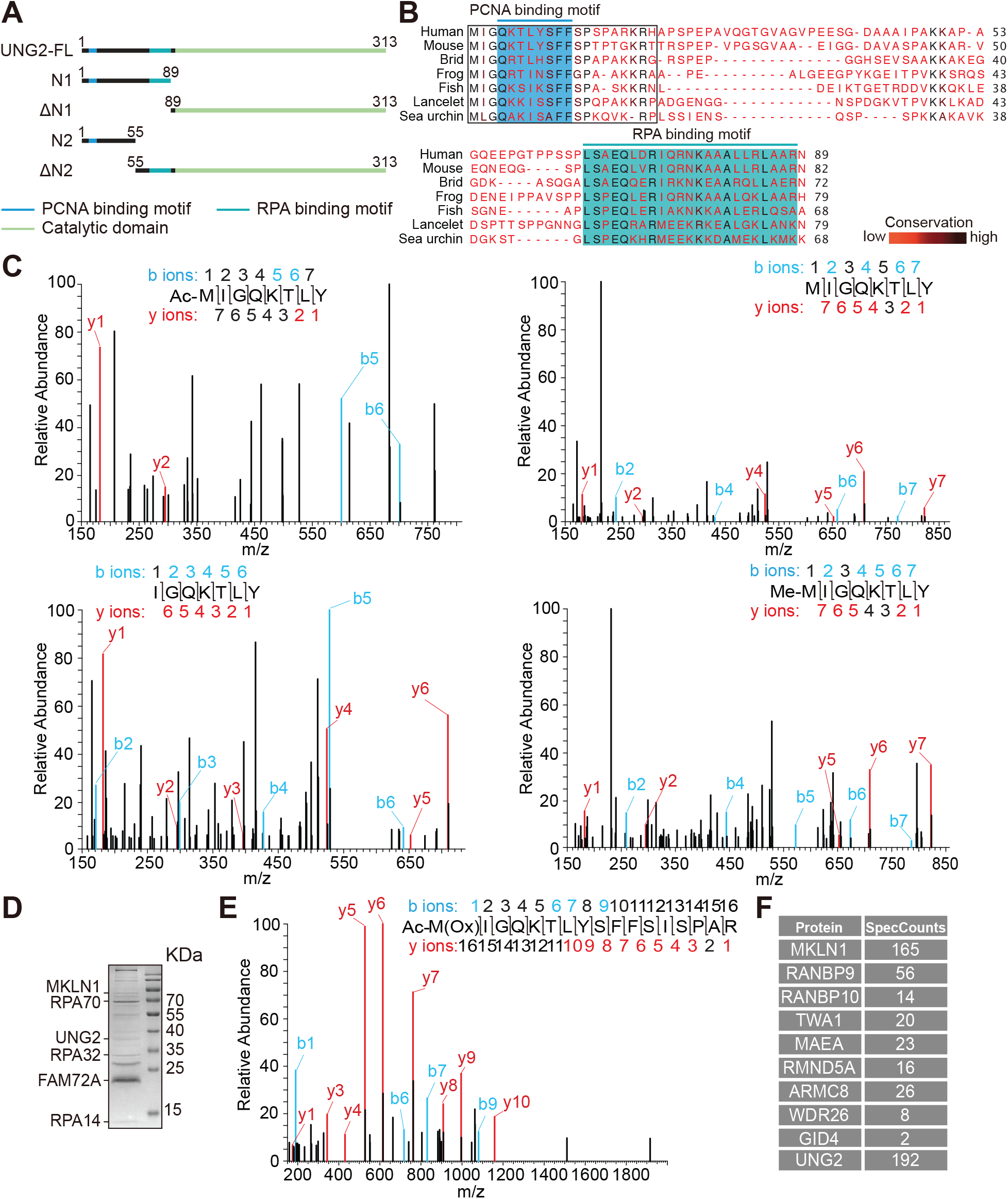
UNG2 interacts with the CTLH^MKLN1-FAM72A^ E3 complex through N-terminal acetylation. (A) Domain organization and construct design of UNG2. (B) Sequence alignment of the N-terminal flexible regions of UNG2 orthologs from human (*Homo sapiens*), mouse (*Mus musculus*), bird (*Taeniopygia guttata*), frog (*Xenopus tropicalis*), fish (*Danio rerio*), lancelet (*Branchiostoma belcheri*), and sea urchin (*Strongylocentrotus purpuratus*). Residues are colored from red to black according to sequence conservation, with red indicating low conservation and black indicating high conservation. The PCNA- and RPA-binding motifs are highlighted in blue and green, respectively. The N-terminal peptide (1-20Aa) used for ITC assays is outlined in black. (C) MS/MS fragmentation spectra of the distinct N-terminal processing products of UNG2 identified from the protein bands indicated by red arrows in Fig. 1B (lanes 6 and 7). (D) Representative SDS-PAGE gel of an affinity pull-down assay using Strep-tagged FAM72A as bait. Proteins captured from HEK293T whole-cell lysates were detected by Coomassie staining, and the identified protein bands are indicated. (E) MS/MS fragmentation spectrum of an N-terminally acetylated peptide from endogenous UNG2 pulled down by FAM72A. The protein band used for MS analysis is indicated in (D). (F) Spectral counts of the CTLH complex components and the substrate UNG2 captured in AP-MS using FAM72A as bait.

**Figure EV2.**
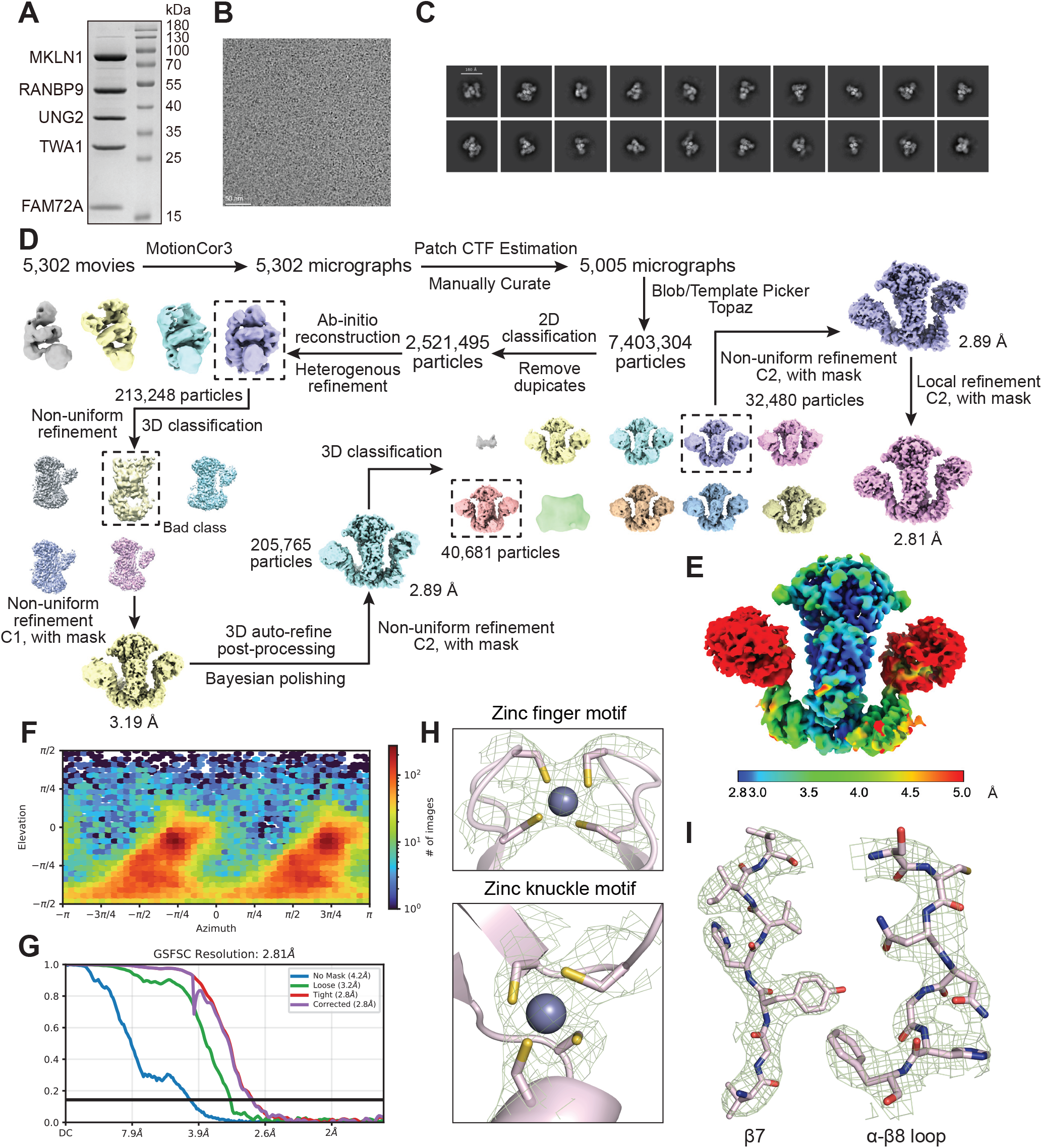
Cryo-EM analysis of the UNG2-FAM72A-MKLN1 complex by single-particle reconstruction. (A) SDS-PAGE analysis of the purified UNG2-FAM72A-MKLN1-RANBP9-TWA1 complex showing equimolar stoichiometry. (B) Representative cryo-EM micrograph of the UNG2-FAM72A-MKLN1-RANBP9-TWA1 complex. Scale bar, 50 nm. (C) Representative 2D class averages from the cryo-EM dataset. Scale bar, 10 nm. (D) Workflow for single-particle cryo-EM analysis of the UNG2-FAM72A-MKLN1-RANBP9-TWA1 complex. (E) Local resolution map of the UNG2-FAM72A-MKLN1 complex, with resolutions ranging from 2.8 to 5.0 Å. (F, G) Particle viewing direction distribution and FSC curves for the final cryo-EM reconstruction. (H, I) Representative cryo-EM density (green mesh) for the zinc finger motif, zinc knuckle motif, β7 strand, and α-β8 loop of FAM72A (pink model). Density maps are displayed in PyMOL at a contour level of 16 σ or 18 σ.

**Figure EV3.**
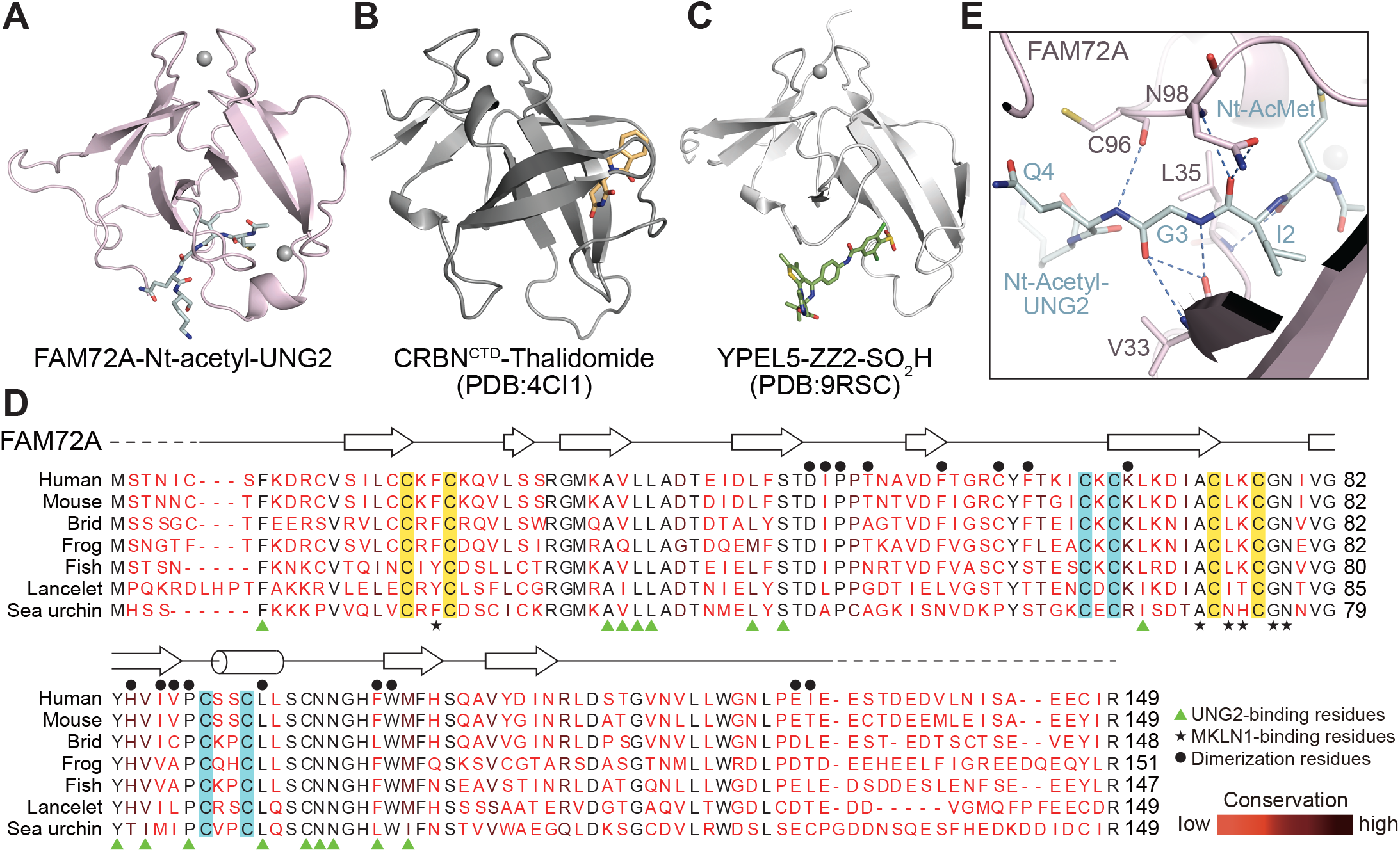
Structural features of FAM72A within the interaction network. (A-C) Comparison of the CULT domain fold adopted by FAM72A, CRBN-CTD, and YPEL5. The three structures are shown as cartoon representations in the same orientation. The bound ligands, the Nt-acetyl-UNG2 peptide, thalidomide, and ZZ2-SO_2_H are shown as teal, yellow, and green sticks, respectively. Zinc ions are shown as gray spheres. (D) Sequence alignment and secondary structure annotation of FAM72A orthologs from human (*Homo sapiens*), mouse (*Mus musculus*), bird (*Taeniopygia guttata*), frog (*Xenopus tropicalis*), fish (*Danio rerio*), lancelet (*Branchiostoma belcheri*), and sea urchin (*Strongylocentrotus purpuratus*). Secondary structure elements are shown as cylinders (α-helices) and arrows (β-strands). Dashed lines indicate regions that are disordered in the solved structure. Residues are colored from red to black according to sequence conservation, with red indicating low conservation and black indicating high conservation. Cysteine residues coordinating zinc ions in the zinc finger and zinc knuckle motifs are highlighted in yellow and blue, respectively. Green arrows and black stars below the alignment indicate residues critical for UNG2 and MKLN1 binding, respectively, while black dots above the alignment denote residues involved in FAM72A dimerization. (E) Interaction network stabilizing the backbone configuration of Nt-Acetyl-UNG2 bound to FAM72A. Nt-Acetyl-UNG2 is shown as teal sticks, and FAM72A is shown as pink ribbon representation. Selected interface residues are displayed as sticks, and blue dashed lines indicate hydrogen bonds.

**Figure EV4.**
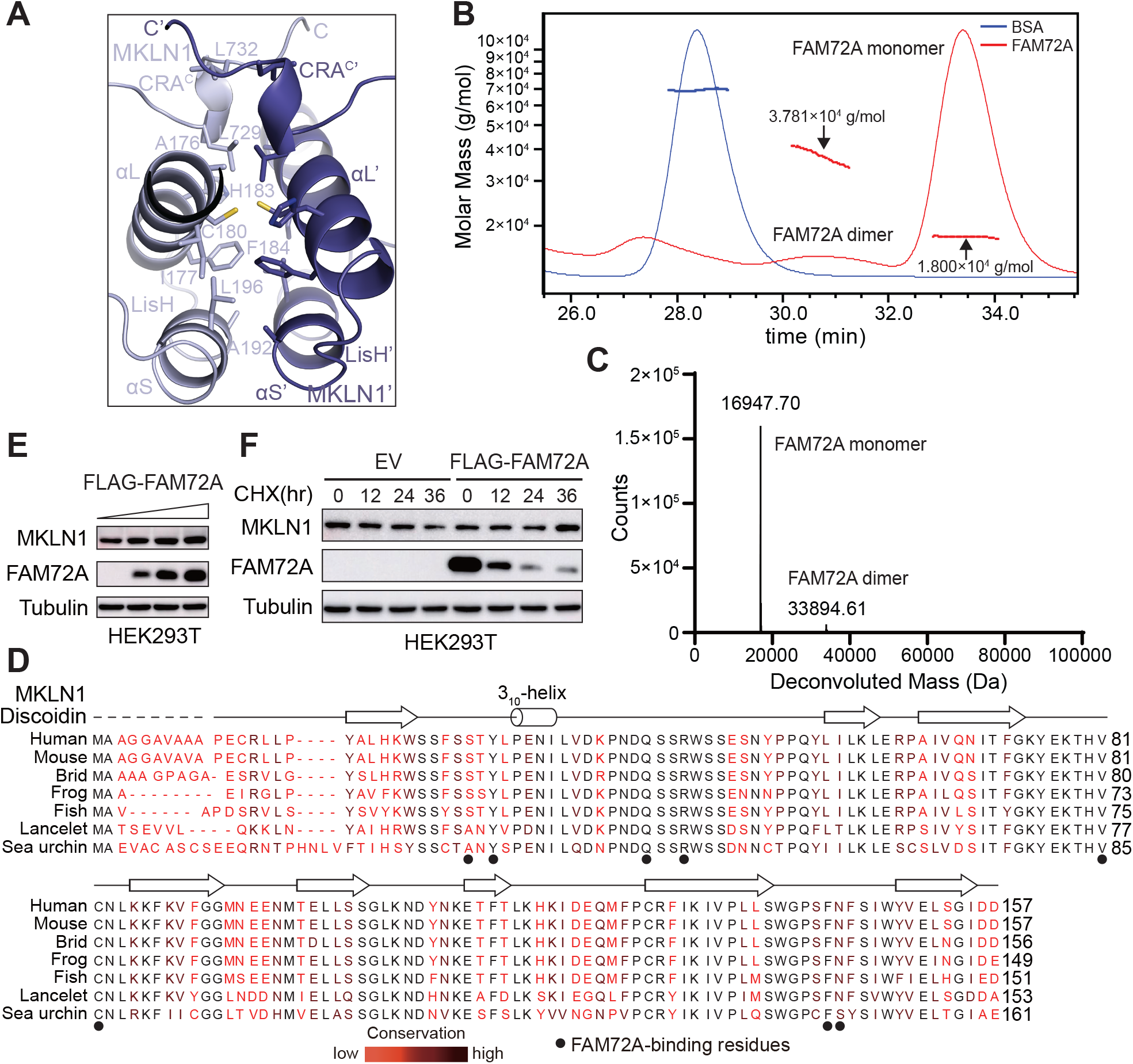
FAM72A is recruited to the CTLH^MKLN1^ E3 complex as a dimer and stabilizes MKLN1 in cells. (A) Hydrophobic dimerization core of MKLN1 formed by the LisH and CRA^C^ domains. MKLN1 is shown as slate blue ribbons, and selected residues at the dimer interface are displayed as sticks. (B, C) SEC-MALS and native mass spectrometry analyses of purified FAM72A. BSA was used as the SEC-MALS standard. The experimentally determined molecular weights are indicated. (D) Sequence alignment and secondary structure annotation of the discoidin domain of MKLN1 orthologs from human (*Homo sapiens*), mouse (*Mus musculus*), bird (*Taeniopygia guttata*), frog (*Xenopus tropicalis*), fish (*Danio rerio*), lancelet (*Branchiostoma belcheri*), and sea urchin (*Strongylocentrotus purpuratus*). Secondary structure elements, including 3_10_-helix and β-strands, are represented as cylinders and arrows, respectively. Dashed lines indicate regions that are disordered in the solved structure. Residues are colored from red to black according to sequence conservation, with red indicating low conservation and black indicating high conservation. Black dots below the alignment denote residues involved in FAM72A binding. (E) Assessment of the steady-state abundance of endogenous MKLN1 in HEK293T cells following transient transfection with increasing amounts of FAM72A plasmid. Whole-cell lysates were analyzed by immunoblotting with the indicated antibodies. (F) Cycloheximide (CHX) chase assays monitoring the stability of endogenous MKLN1. HEK293T cells were transiently transfected with the indicated plasmids. 24 h after transfection, cells were treated with CHX for 0-36 h. Whole-cell lysates collected at the indicated time points were analyzed by immunoblotting with the indicated antibodies. EV, empty vector.

**Figure EV5.**
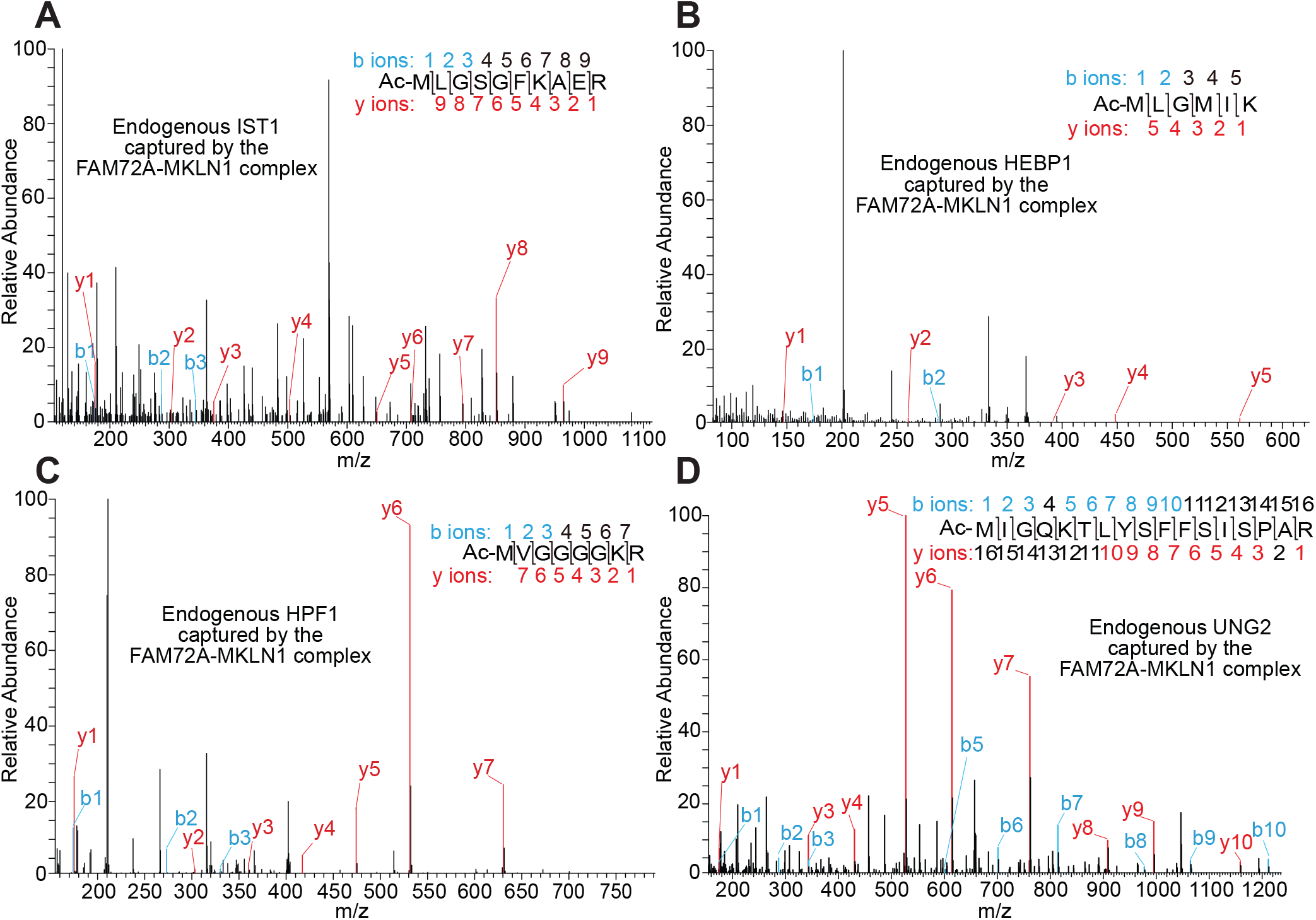
N-terminal acetylation of the neo-substrates recognized by FAM72A. (A-D) MS/MS fragmentation spectra of N-terminally acetylated peptides from candidate substrates captured by AP-MS using the FAM72A-MKLN1 complex as bait. All the observed *m/z* values of the b1 ions are consistent with N-terminal acetylation of the methionine residues.

**Figure EV6.**
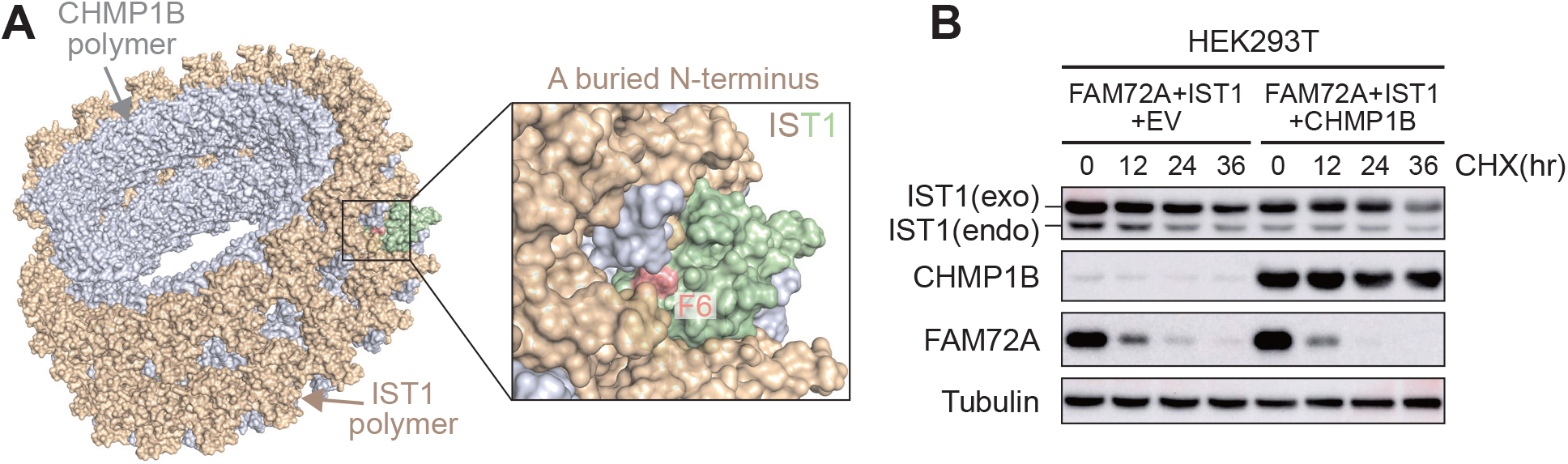
Context-dependent IST1 degradation through the AcM-[ILV]-G/N-degron pathway. (A) Double-stranded helical filament assembly formed by IST1 and CHMP1B (PDB: 6TZ4). All proteins are shown as surface representations, with IST1 colored in wheat and CHMP1B in light gray. A representative IST1 monomer is shown in pale green, with its first visible N-terminal residue (F6) highlighted as salmon sticks and surface. The enlarged close-up view reveals that F6 is deeply buried within a pocket formed by IST1 and CHMP1B, suggesting that the N-terminal degron of IST1 is likely inaccessible to the substrate receptor FAM72A in this assembly context. (B) Cycloheximide (CHX) chase assays monitoring the cellular stability of IST1. HEK293T cells were transiently transfected with the indicated plasmids. 24 h after transfection, cells were treated with CHX for 0-36 h. Whole-cell lysates collected at the indicated time points were analyzed by immunoblotting with the indicated antibodies. EV, empty vector.

**Table EV1.** Cryo-EM data collection, refinement and validation statistics

|  | UNG2-FAM72A-MKLN1<br>(EMD-82509)<br>(PDB 44CV) |
| --- | --- |
| **Data collection and processing** |  |
| Magnification | 105,000 |
| Voltage (kV) | 300 |
| Electron exposure (e-/Å<sup>2</sup>) | 60 |
| Defocus range (µm) | -1.0 to -2.0 |
| Pixel size (Å) | 0.821 (physical) |
| Symmetry imposed | C2 |
| Initial particle images (no.) | 2,521,495 |
| Final particle images (no.) | 73,161 |
| Resolution at 0.143 FSC threshold (Å) | 2.81 |
| Map resolution range (Å) | 2.8-5.5 |
| **Refinement** |  |
| Initial model | AlphaFold 3 |
| Model resolution (Å) | 2.8 |
| FSC threshold | 0.143 |
| Map sharpening *B* factor (Å<sup>2</sup>) | -60.6 |
| Model composition |  |
| Non-hydrogen atoms | 6918 |
| Protein residues | 848 |
| Zn<sup>2+</sup> | 4 |
| *B* factors (Å<sup>2</sup>) |  |
| Protein | 158.16 |
| Zn<sup>2+</sup> | 194.64 |
| R.m.s. deviations |  |
| Bond lengths (Å) | 0.003 |
| Bond angles (°) | 0.661 |
| Validation |  |
| MolProbity score | 1.75 |
| Clashscore | 7.11 |
| Poor rotamers (%) | 1.02 |
| Ramachandran plot |  |
| Favored (%) | 94.93 |
| Allowed (%) | 5.07 |
| Disallowed (%) | 0 |

